# Morphological transformation in *Helicobacter pylori* is a dynamic process leading to two types of coccoid

**DOI:** 10.64898/2026.01.13.698397

**Authors:** Thimoro Cheng, Phuong-Y Mai, Abimbola Feyisara Adedeji-Olulana, Olivia E. R. Smith, Richard Wheeler, Joel Berry, Samia Hicham, Olivier Danot, Simon Foster, Tanmay A.M. Bharat, Jamie K. Hobbs, Ivo Gomperts Boneca

**Affiliations:** Institut Pasteur, Université Paris Cite, INSERM U1306, CNRS UMR6047, Unité Biologie et génétique de la paroi bactérienne, Paris, France; School of Mathematical and Physical Sciences, University of Sheffield, Sheffield, UK; Structural Studies Division, MRC Laboratory of Molecular Biology, Cambridge CB2 0QH, United Kingdom; School of Biosciences, University of Sheffield, Sheffield, UK; The Florey Institute, University of Sheffield, Sheffield, UK

**Author notes:** Author Contributions: T.C. and I.G.B designed the study, T.C., P-Y.M., A. F. A-O., O.E.R.S, R.W., J.B., S.H. and O.D. performed the experiments, T.C., O.D and I.G.B wrote the article, S.F., T.A.M.B, J.K.H. and I.G.B. supervised different aspects of the project. **Competing Interest Statement:** The authors declare no competing interests.

**Keywords:** Peptidoglycan, Helicobacter pylori, coccoid, cell surface, cell shape, microscopy

## Abstract

The helical shape of *Helicobacter pylori* is crucial for successful colonization of the human stomach. However, this pathogen can shapeshift into another form termed the “coccoid form” with a spherical shape, through a mechanism that remains elusive. Here, by a combination of fluorescence microscopy using fluorescent D-aminoacids, cryoelectron and atom force microscopy, we explored the dynamics of coccoid formation in *H. pylori* through interrogation of the peptidoglycan layer. Contrary to the widely held hypothesis, we showed that helical-rod *H. pylori* transformed into a coccoid without transiting through a U-form. We show that U-forms, characterized by a U-shaped peptidoglycan with enlarged periplasmic space, altered genetic material, and red autofluorescence, are the output of a parallel pathway, which, unlike the coccoid pathway, is independent of the HdpA/Csd3 peptidoglycan endopeptidase. Coccoid formation occurred along a rigid timeline, by bulging of the cytoplasmic membrane through a peptidoglycan crack, resulting in a spheroplast-like structure with the peptidoglycan stacked into a thick layer near the original cell poles. Resistance of that structure against lysis likely involves a switch in metabolic profile reminiscent of bacteria in dormancy, with a notable accumulation of lysophospholipids, demonstrated in this work. Altogether, the ultrastructure and properties of *H. pylori* coccoids evidenced here are compatible with a role of this form in relapse after antibiotic treatment.

**SIGNIFICANCE STATEMENT:** Upon prolonged growth, stomach pathogen *Helicobacter pylori* undergoes a morphological change from a helical to a spherical form called coccoid, which may be involved in bacterial persistence and immune evasion. The pathway leading to this form, as well as its precise architecture, were unclear. In this work, we show by different microscopy techniques that the transition to coccoid does not involve a U-shaped intermediate as proposed before, but is triggered by progressive thinning, due to the activity of endopeptidase HdpA/Csd3, of the peptidoglycan meshwork that normally protects the cell, eventually leading to rupture. Because of the hole thus created, peptidoglycan can no longer contain the osmotic pressure in the cytoplasm, which leaks out within a membrane bulge, to eventually give rise to a kind of sphaeroplast, expected to be insensitive to cell wall-targeted antibiotics.

## INTRODUCTION

*Helicobacter pylori* is a carcinogenic bacterium responsible for causing chronic gastritis, peptic ulcers, and gastric cancer (1–3). For millennia, this organism has been colonizing and co-evolving with its only host, humans, and is currently found in half of the world’s population (3). To successfully colonize, *H. pylori* must survive the harsh acidic environment of the human stomach and move towards its preferred niche, the basal layer of gastric epithelium. *H. pylori’s* ability to neutralize pH, its flagellar-driven motility, and capacity to adhere to host cells as well as its virulence factors, CagA and VacA, have all been shown to be crucial in this process (4). Recently, the helical shape of *H. pylori* has been proposed as a virulence factor since this organism was found to fail to colonize when its peculiar shape was compromised (5, 6). This helical shape was postulated to allow these bacteria to bore into the mucus layer in a corkscrew-like motion and to efficiently swim to escape clearance by mucus sloughing and peristalsis. Fascinatingly, *H. pylori* has been observed to live a dimorphic lifestyle and can transform from helical to spherical cells termed ‘coccoid form’. This latter form has been demonstrated to possess a distinctive peptidoglycan composition profile likely to promote immune evasion from NOD1, an innate immune receptor recognizing peptidoglycan (PGN) and a major responder in *H. pylori* infection (7, 8). Additionally, the coccoid form has been proposed to be in the viable-but-non-culturable (VNBC) state of dormancy and to participate in antibiotic tolerance and transmission (1).

Bacterial cell shape is largely determined by the PGN or cell wall, a component of the bacterial cell envelope. It is a network consisting of glycan chains of repeating disaccharide, *N-*acetylglucosamine (NAG) ß1,4 *N-*acetylmuramic acid (NAM), crosslinked by peptide stems (9). The PGN structure is the outermost layer of the cell envelope in Gram-positive or monoderm bacteria that lack a surface layer (S-layer) but is sandwiched in between the inner and outer membranes in diderm species such as *H. pylori.* While peptidoglycan sturdiness is essential to withstand the turgor pressure of the cytosol, PGN is also a highly dynamic structure that is constantly expanded and remodeled to accommodate cell growth and division. Model rod-shaped bacteria like *Bacillus subtilis* and *Escherichia coli* achieve their morphology by the functions of two separate PGN-remodeling complexes, the divisome and the elongasome (9). Although the *H. pylori* genome encodes most of the components of the elongasome, it harbors a reduced set of PGN biosynthetic genes in comparison with other bacteria (10). Previous work in our laboratory and by others identified and characterized several cell shape determinants in *H. pylori* including Csd1-2, HdpA/Csd3, Csd4, Csd5, Csd6, CcmA, MreB, and AmiA (5, 8, 11–15). However, how *H. pylori* transitions towards the coccoid shape and what the genetic and environmental determinants of this process are, remains unclear. In particular, whether U-shaped cells, which appear at the same time as coccoid cells (16–18), represent an intermediate in the transition or the endpoint of a parallel morphogenetic pathway is unknown.

Here, we established strain N6 of *H. pylori* to be an ideal strain for studying coccoid formation. Using timelapse fluorescence imaging, we precisely describe the steps leading from helical rod to either spherical coccoid or U-form. Characterization of the kinetics of this process identified a transition period during which the metabolic state of the bacteria switched and lysophospholipids (LPLs) accumulated. Ultrastructural studies with cryo-electron (cryo-EM) and atomic force microscopy revealed that the two forms possess clearly distinct cell envelope structures. U-forms further differed from coccoids by their genetic content, showing a characteristic autofluorescence, and their appearance was independent of the action of HdpA/Csd3. Altogether, these findings deepen our understanding of the coccoid form of *H. pylori* and suggest the existence of two different subpopulations with respect to their fate in stationary phase.

## RESULTS

### Strain N6 as a model for studying coccoid formation in *H. pylori*

*H. pylori* is well-known for its high genotypic and phenotypic variations among strains. To determine which strain is the most suitable to analyze coccoid formation dynamics, we examined this process in 8 strains of *H. pylori* commonly used in laboratory research (26695, B128, G27, J99, N6, P12, SS1, X47, Table S1) by phase-contrast microscopy. All 8 strains were observed to enter coccoid transition at the beginning of the stationary phase and remained in that state until the end of the experiment (Fig. S1A, S2). We then compared the 8 strains based on 4 parameters: rod-to-coccoid size ratio, aggregation tendency, transition kinetics, and genetic tractability (Fig. S1B and Table S2). According to these parameters, N6 seemed the most appropriate strain for this study: it had a high rod-to-coccoid size ratio (2.3), a low aggregation tendency, ideal transition kinetics (transition at ∼40 hours), and it easily takes up plasmid DNA through natural transformation.

### Kinetics of coccoid formation in strain N6

We analyzed the growth kinetics and culturability of strain N6 in liquid culture over a period of 2 weeks. In this strain, stationary phase began around T35-T40 and culturability strongly decreased at T90 to reach 0 at T130-T150 in our conditions (Fig. 1A, B). Using microscopy imaging and the machine learning tool ilastik, we tracked the proportion of the helical-rod and the coccoid form over the span of its culturable period (140 hours). Analysis on over 70,000 cells confirmed that coccoid formation in strain N6 initiated in the beginning of the stationary phase with most of the transition occurring between T45 and T65 (Fig. 1C, D).

**Figure 1:**
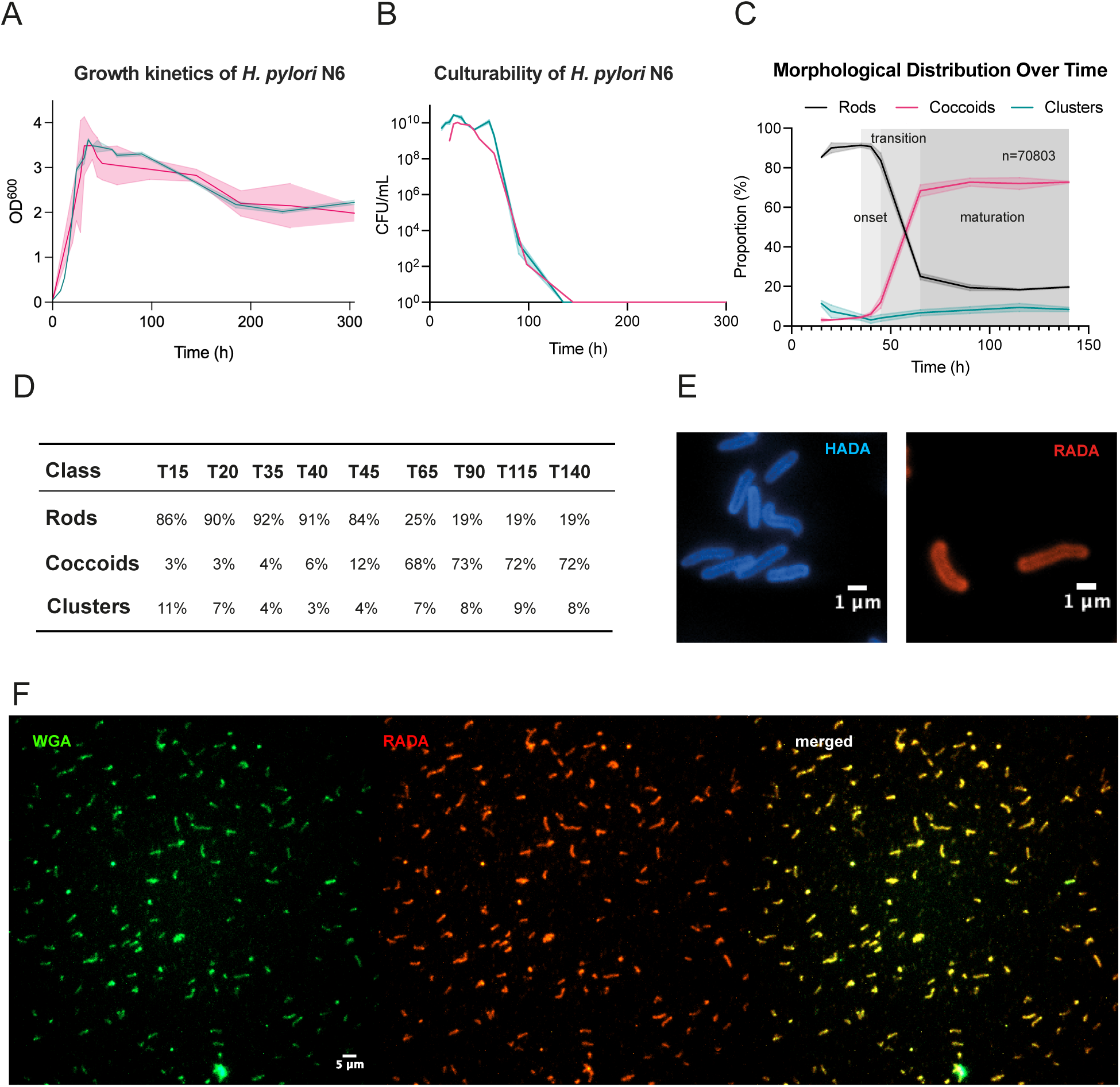
Kinetics of the *H. pylori* morphogenesis. A-B) Growth kinetics and culturability. Bacteria were grown in replicates for a period of 2 weeks in liquid cultures in two separate experiments and aliquots were taken at various timepoints for OD600 measurement and CFU assay on plates. Data was truncated to 300 hours and visualized with GraphPad. C) Morphological distribution of *H. pylori* strain N6 over time. Bacteria were grown in replicates for a period of 140 hours in liquid cultures and fixed at various timepoints for imaging on phase contrast. Data was pooled from 3 independent experiments. Morphological analysis was conducted in ilastik and results were imported into GraphPad for visualization. D. Morphological distribution of *H. pylori* over time. E-F) FDAAs in *H. pylori*. E) Fluorescence images of *H. pylori* N6 stained with HADA (blue) and RADA (red) taken at T20 of growth in liquid culture. F) Fluorescence images of RADA-incorporated *H. pylori* sacculi isolated at T20 of growth and stained with wheat germ agglutinin (WGA).

We segmented this morphological transformation into three distinct phases: onset, transition, and maturation (Fig. 1C, D). The onset phase marks the first increase in the coccoid proportion and initiates at the beginning of the stationary phase at round T35-T45. In this phase, both cell elongation and division have ceased. Subsequently, the transition phase occurs and persists for about 4-5 generation times, spanning between T45 and T65. During this period, the majority of the morphological transformation takes place, and the coccoid proportion jumps drastically from around 12% to almost 70%. At the maturation phase coccoid formation becomes slow but detectable until T90.

### FDAAs as molecular probes for tracking the *H. pylori* PGN

Since peptidoglycan is the major determinant of bacterial cell shape, we decided to use fluorescent D-amino acids (FDAAs) to visualize PGN remodeling in *H. pylori.* FDAAs have been shown to be incorporated into the peptide stems of PGN at the 4^th^ or 5^th^ position, by L,D-transpeptidases (TPases) and D,D-TPases, respectively (19). They can also be cleaved off by D,D-carboxypeptidases (CPases) such as PBP4, PBP5 or DacA in *E. coli* (19). Simply growing wildtype N6 *H. pylori* in the presence of one of two different FDAAs, RADA or HADA, was sufficient to fluorescently label all cells, indicating that the outer membrane is permeable to these compounds (Fig. 1E). RADA was covalently incorporated into the PGN since purified RADA-labeled sacculi stained with wheat germ agglutinin (WGA) showed colocalization of the two fluorescence signals (Fig. 1F).

*H. pylori* PGN is rich in pentapeptides and possesses only 4-3 crosslinks (7, 8). FDAAs are therefore expected to be incorporated into the 5^th^ position of the peptide stem by the three PBPs of *H. pylori*: PBP1, PBP2, and PBP3, as described by Kuru *et al.* (19). To test this prediction, we conducted pulse-chase experiments with RADA and HADA in the presence of aztreonam and meropenem that preferentially inhibit PBP3 and PBP2 of this species, respectively. We first labeled *H. pylori* cells with HADA, then treated them with either aztreonam or meropenem and chased with RADA for 20 minutes (8% of a generation time). By fluorescence imaging and image analysis, RADA was found to be incorporated only at the septum (and the new pole after division) when the bacteria were treated with meropenem indicative of the PBP3 (divisome) activity (Fig. S3). Conversely, it was selectively incorporated into the lateral body of the filamented cells obtained when treated with aztreonam, reflecting the PBP2 activity (elongasome). This suggested that PBP2 and PBP3 are involved in FDAAs incorporation into the PGN of *H. pylori,* and consequently that this incorporation occurs at the 5^th^ position of the peptide stem.

### FDAA labeling revealed coccoid form with the two poles close to each other

To inspect the cell wall architecture of the coccoid form, we used RADA and HADA . We cultured the bacteria in the presence of either probe until late stationary phase to generate a mainly coccoid population and observed the labeling pattern in this form by fluorescence imaging. While the FDAA was homogeneously distributed along the lateral body of helical-rod *H. pylori,* it localized on one side of the coccoid form rather than around the entire spherical sac (Fig. 2A). This suggested that the PGN in the coccoid form was highly remodeled. By contrast, the lipophilic dye FM4-64x stained the entire outline of the coccoid cells, suggesting that the membranes were intact (Fig. 2B). Hoechst 3342 labelling indicated the presence of DNA, which localized within the spherical sac surrounded by lipid membranes and on one side of the PGN as shown by co-staining with FM4-64x and RADA , respectively (Fig. 2B). These data indicated that the coccoid form of *H. pylori* comprises a nucleoid surrounded by lipid membranes, presenting a highly remodeled PGN with the two poles close to each other.

**Figure 2:**
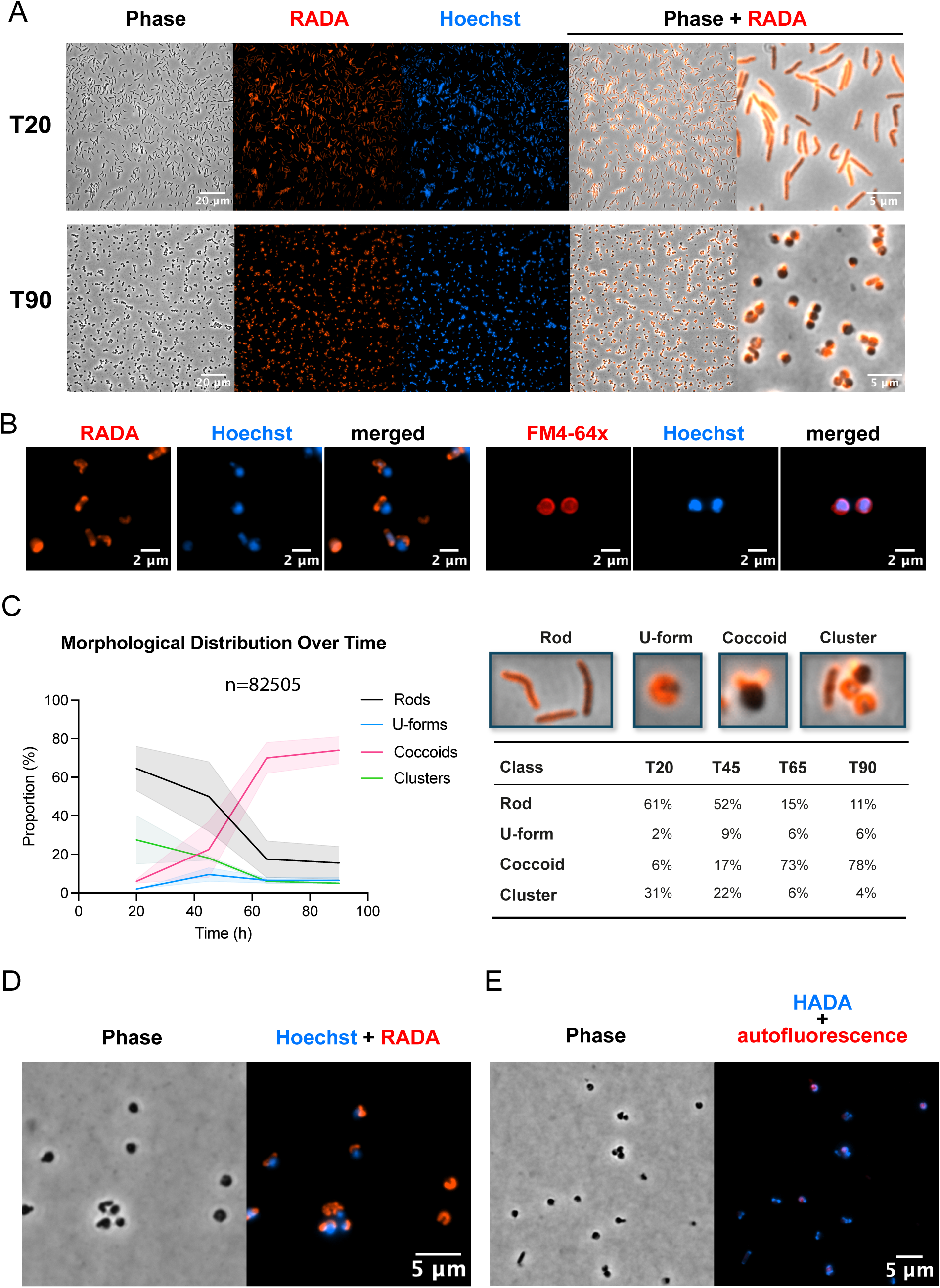
*H. pylori* morphogenesis during the transition to the coccoid form. A) Helical-rod (top panel) and coccoid *H. pylori* (bottom panel) grown in the presence of RADA (red) and stained with Hoechst 3342 (blue) imaged after 20 hours and 90 hours of growth, respectively. B) Overlay fluorescence images of *H. pylori* coccoid labeled with (a) WGA (red) and HADA (blue); (b) FM4-64fx (red) and HADA (blue); (c) RADA (red) and Hoechst (blue); and (d) FM4-64x (red) and Hoechst (blue). C) Left: morphological distribution of *H. pylori* during the transformation. Bacterial cells were cultured in the presence of RADA in replicates and imaged at 4 different timepoints. Morphological analysis was conducted with ilastik and results were pooled from 3 independent experiments and visualized using GraphPad. Right: Representative images of the four morphological classes and relative proportions at the 4 timepoints. D) U-form in *H. pylori. H. pylori* cells at T90 labeled with RADA (red) and Hoechst 3342 (blue) show U-forms lacking DNA. E) *H. pylori* cells at T65 labeled with HADA (blue) and imaged for autofluorescence signal (red).

### The U-form is not a coccoid intermediate but belongs to a parallel morphogenetic pathway

Many studies reported observation of a U-shaped intermediate as *H. pylori* transforms from helical-rod into coccoid (16–18). Even if no such form could be distinguished by ilastik in the phase-contrast analysis (Fig. 1), we observed in the FDAA labelling experiments that some cells with a spherical morphology in phase contrast had a U-shaped or donut-like fluorescence signal (Fig.2C). These cells will be called U-forms in the rest of the article. To understand the role of the U-forms in the coccoid transition, we quantified four morphotypes along growth (rod cells, U-forms, coccoid and clusters) using ilastik, this time from fluorescence microscopy images. The proportion of the coccoid form jumped dramatically during the transition phase between T45 and T65, and only slightly increased between T65 and T90. The proportion of the U-forms increased slightly at T45 and then remained relatively constant until T90 (Fig. 2C), where a decrease would have been expected in the hypothesis of an intermediate form. Moreover, we noticed that these U-forms frequently displayed features distinguishing them from coccoid cells: they generally did not stain with Hoechst, DAPI or SYTO-13, three dyes used for DNA-staining (Fig. S4), and they emitted autofluorescence in the red spectrum at a higher intensity compared to other forms (Fig. 2D, E). Altogether, these results casted serious doubt on the U-shaped intermediate hypothesis.

To visualize the morphological intermediates on the pathway to the coccoid form at the single-cell level, we conducted a time-lapse imaging of N6 live cells pre-labeled with RADA. The bacteria were first grown in liquid culture supplemented with RADA until stationary phase (T45), then transferred onto BHI-agarose pads for imaging in a growth-permissive environment. All *H. pylori* cells observed transformed into coccoids, not via a U-shaped intermediate, but rather through a swelling or bulging mechanism (Fig. 3A, Videos S1-2).

**Figure 3:**
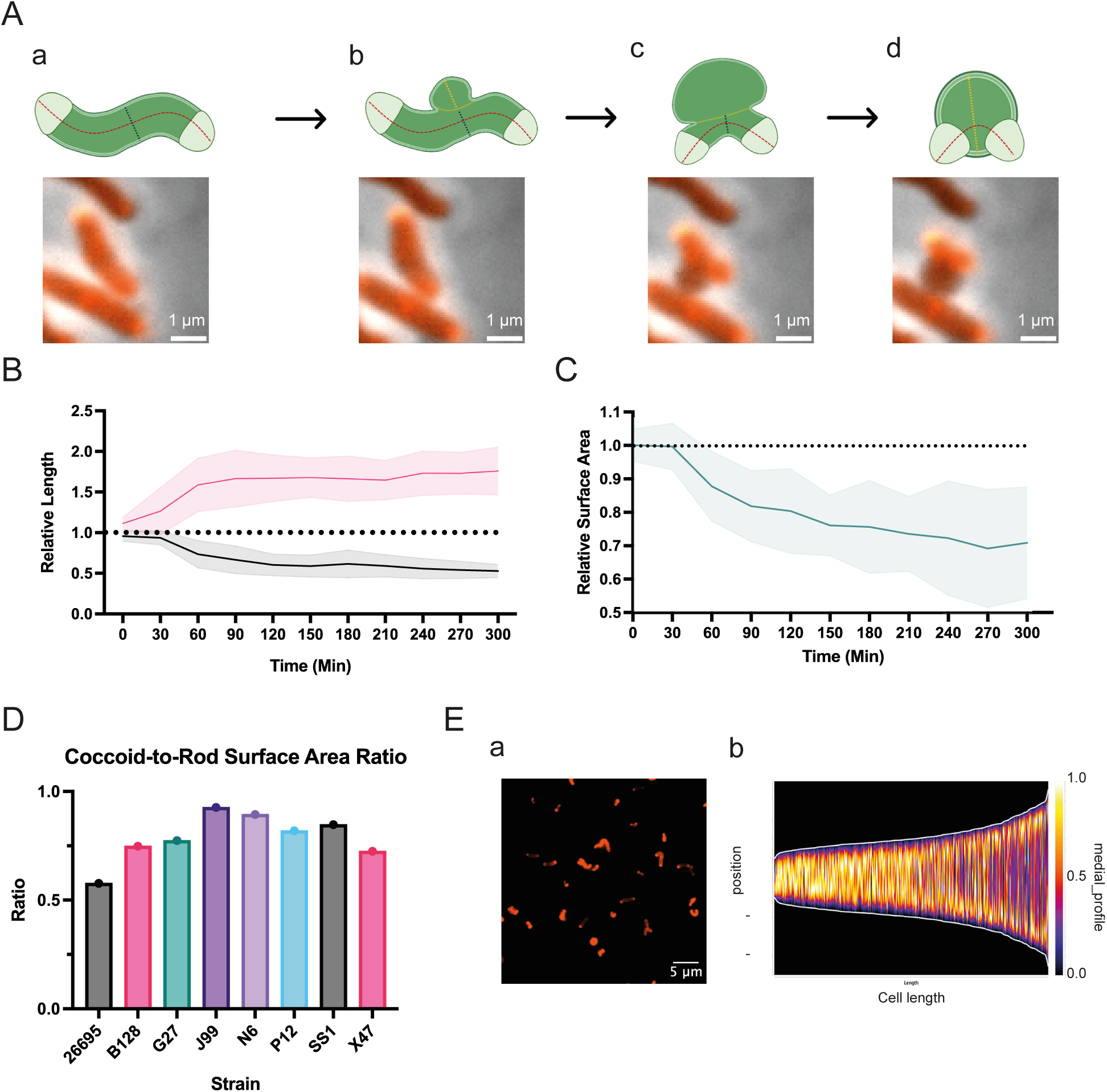
Cell wall architectural dynamics during morphogenesis. A) Cartoon representation of the cell shape dynamics during the transformation with snapshots of *H. pylori* rod transformed into coccoid. Each image is an overlay of RADA signal (orange) and phase contrast (black), cropped from time-lapse imaging data at 30-min intervals. B) Relative change in cell length and bulge size during the morphological transformation measured from the time-lapse imaging data. C) Relative change in surface area during the morphological transformation. Surface area was calculated using rod length and bulge diameter measured from the time-lapse data and normalized by the value of the first timepoint. D) Coccoid-to-Rod surface area ratio calculated at T10 and T135 of *H. pylori* strains with the assumption of straight rod (modeled as a cylinder closed by two hemispheres) and perfect sphere, respectively. E) Dynamics of RADA intensity during the transformation. (a) Fluorescence image of *H. pylori* cells at T65 grown in the presence of RADA (red) and (b) demograph of RADA signal measured along the lateral body of *H. pylori* cells captured at T65 sorted by length from shortest (left) to longest (right).

### Cell shape dynamics during the morphological transformation

According to the time-lapse imaging data, initiation of the bulging process occurred at random sites along the lateral body of the bacterial cells. During the transition process (Fig. 3A), the length of the cells decreased by half while their width at the bulge (bulge diameter) increased by 1.6 compared to the initial width of the rod (Fig 3B, C). Most of these changes took place in a span of 60-90 minutes after initiation.

Spheres are the shapes with the lowest surface-to-volume ratio. Using the length and bulge diameter measured, we calculated the surface area of 11 bacterial cells as they transformed from rod to coccoid and found it to decrease by 25% on average (Fig. 3C). Similarly, an at least 10% decrease in surface area was observed between rod cells of different strains at T10 (log phase) and matured coccoid cells at T135 (late stationary phase) (Fig. 3D). The 25% to 10% difference for N6 was possibly due to the fixation of cells in the latter experiment.

Additionally, we observed an inverse relationship between the FDAA intensity and the rod length during the transformation. The fluorescence signal in the lateral body of the helical rod intensified as its length decreased over time. Thus, the RADA intensity around the contour of cells from T65 (end of transition phase) consisted of mixed phenotypes:rod cells with bright poles but relatively lower RADA signal in the lateral body and shorter rods or coccoid cells with higher intensity all around. A demographic profile of the RADA signal indeed showed accumulation of the signal at the poles of longer cells but homogenous distribution in shorter cells (Fig. 3E). The dramatic reduction in cell length observed after the morphological transformation along with the intensification of the FDAA signal suggests that the PGN layer of *H. pylori* undergoes stacking during this process.

### The spherical sac of the coccoid form is devoid of PGN

The morphological transformation is reminiscent of that of beta lactam-treated *E. coli* in which the PGN layer breaks with the intracellular content blebbing out due to the turgor pressure (22, 23). We speculated this could also be the case for coccoid *H. pylori*. To test whether the absence of FDAA labelling of the spherical sac of the coccoid form (Fig. 2A, C) reflects trimmed peptide stems or a total absence of PGN, we extracted RADA-labeled sacculi from the coccoid form (T90) and used wheat germ agglutinin (WGA) to stain the PGN GlcNac moieties. WGA was found to exclusively colocalize with RADA indicating that the spherical sac of the coccoid form is not surrounded by PGN (Fig. S5A).

To analyze the PGN of coccoid cells (T90) in more detail, we used atomic force microscopy (AFM). Consistent with the results above, in air AFM micrographs revealed only rod-shaped, but no spherical sacculi in this mainly coccoid population. Comparison with sacculi from T20 rod cells revealed only minor differences such as a thinner PGN in coccoid-derived sacculi, with a slightly reduced and more variable width (Fig. S5B, C). However, in-liquid AFM showed that the PGN nanoscale architecture of coccoid-and rod-derived sacculi are different. Instead of the regular meshwork of rod-derived sacculi, coccoid-derived sacculi displayed deep transversal cracks suggesting a general weakening of the cross-linking of the structure (Fig. S5D, right-hand side).

To further analyze the cell envelope architecture, we performed ultrastructural studies on *H. pylori* cells captured at T20, T45, and T90 using cryo-EM. We observed very distinctive architectures at different time-points (Fig. S6A) most likely corresponding to the different morphotypes identified by optical microscopy (Fig. S6B). Rod cells (T20) were observed to possess the typical double-membraned cell envelope associated with *H. pylori*. At T45 and T90, we observed cells appearing as bulging rods (T45 top), U-forms (T45 and T90, bottom) and coccoid forms (T90, top). The outer membrane of all these forms appeared intact but locally detached from the inner membrane, leading to an enlarged periplasm.

In the U-form, this enlarged periplasm separated the U-shaped PGN encasing the inner membrane from the spheroidal outer membrane. Cryo-electron tomography (cryo-ET) studies of whole cells were consistent with the cryo-EM two-dimensional images, but the spherical shape and resulting thickness of the coccoid cells prevented the visualization of the very thin *H. pylori* PGN layer in tomograms (Videos S3-S4).

### HdpA is implicated in the formation of coccoid but not U-form

Of the four PGN peptidases identified in *H. pylori,* only HdpA is known to be implicated in coccoid formation. Mutants of *csd1, csd4, and csd6* all undergo the morphological transformation as the wildtype while this process was shown to be delayed in the Δ*hdpA* mutant in *H. pylori* G27 background (5, 11, 13). To quantitatively analyze the effect of HdpA on U-form and coccoid formation, we conducted fluorescence imaging on an N6 Δ*hdpA* mutant labelled with HADA as previously done for the wt. Quantitative analysis on microscopy images of over 53,000 cells showed that the coccoid form represented around 3% of the population, a dramatic difference with the wt at the same time point (around 70%) (Fig. 4A). The proportion of the U-form was similar to that observed in the wild type (around 6% at T90), suggesting that its formation was independent of HdpA/Csd3. Red autofluorescent cells and U-forms amounted to similar proportions except at the last timepoint (8% U-form vs 16% autofluorescence) (Fig. 4A), possibly due to incompletely matured U-forms. Similar results were obtained with with 5% v/v hydrogen peroxide induced coccoids (24) (Fig. 4B), confirming the distinctive role of HdpA in the formation of the coccoid and U-form.

**Figure 4:**
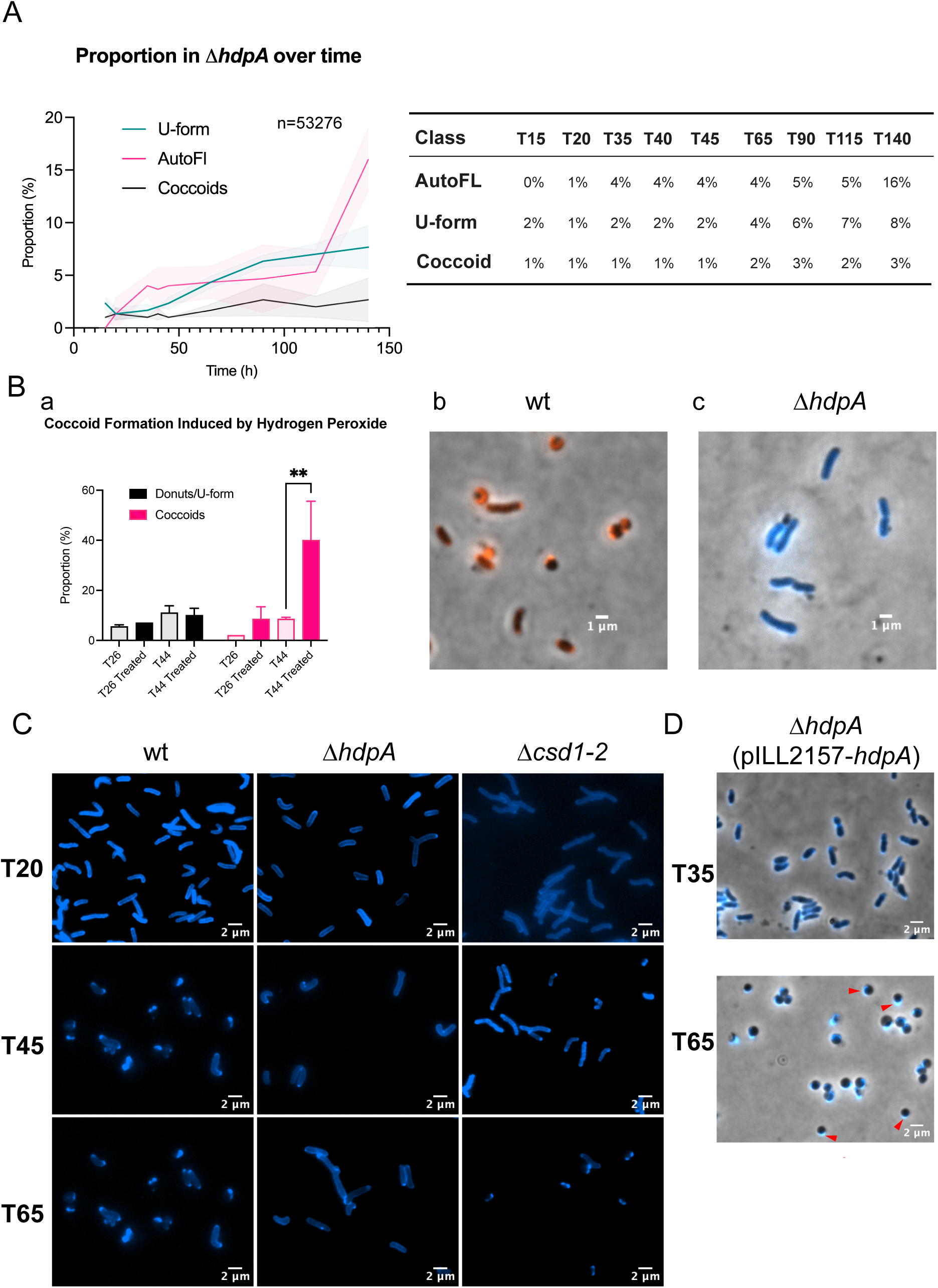
Effects of PGN hydrolases on coccoid formation in *H. pylori.* A) Time-course morphological distribution in *ΔhdpA* mutant highlighting the proportion of the U-form, coccoid form, and autofluorescent cells (AutoFl) over a period of 140 hours. The table displays the values used for the plot. B) Effect of hydrogen peroxide on coccoid formation in wildtype *H. pylori* and *ΔhdpA* mutant. (a) Distribution of the coccoid and the U-form in wildtype strain at T26 and T44 with or without hydrogen peroxide treatment. Bacteria were grown in the presence of RADA (red) or HADA (blue) and 5% hydrogen peroxide was introduced at T20 of growth. Aliquots were taken at T26 (6h post induction) and T44 (24h post induction) for imaging with phase contrast and fluorescence. (b-c) Fluorescence and overlayed images of representative cells at 24h post induction in wildtype with RADA (b) and Δ*hdpA* mutant with HADA (c). C) Fluorescence images of *H. pylori* cells in wildtype strain (a), *ΔhdpA* (b), and *Δcsd1-2* hydrolase mutant (c) at T20, T45, and T65. D) Fluorescence image of *H. pylori* recombinant strain expressing HdpA from an IPTG-inducible plasmid grown in the presence of IPTG and HADA (blue) at T35 and T65. Crescent labelling is indicated by red arrows.

The lower fluorescence signal in the lateral PGN observed in many cells could be due either to differential incorporation of the FDAA along the cell or to a D,D-carboxypeptidase activity specific of the lateral PGN. Since HdpA is the only D,D-carboxypeptidase identified in *H. pylori*, we examined the FDAA labelling pattern in the *ΔhdpA* context. Strikingly, the Δ*hdpA* mutant did not display the lateral signal deficit observed in the wild-type (or in a Δ*csd1-csd2* mutant knocking down the other *H. pylori* D,D-endopeptidase gene), but retained a relatively homogenous pattern (Fig. 4C). This strongly suggests that the low lateral signal is due to removal of the C-terminal, FDAA-conjugated D-Ala by HdpA, and that HdpA activity is largely excluded from the poles. To understand why, we constructed a recombinant strain of *H. pylori* to express HdpA-mScarlet from the *hdpA* locus on the chromosome and conducted a pulse-chase labeling experiment using HADA. The bacteria were first grown in the presence of HADA (long-pulse), and were then transferred into a fresh medium devoid of the probe and were allowed to grow for half a generation time (chase). Image analysis revealed the loss of HADA to coincide with the HdpA-mScarlet signal along the lateral body (Fig. S7) indicating that the subcellular localization of HdpA activity is due to the localization of the protein itself. The main activity of HdpA (D,D-endopeptidase) is therefore also likely to follow that pattern. Altogether these results suggest that HdpA acts in the coccoid transition through remodeling of the lateral PGN, most likely by hydrolyzing the cross-links between glycan strands of lateral PGN.

We previously showed that overexpression of HdpA induced spherical cells in *H. pylori* (12). Because of their shape and of the role of HdpA in coccoid formation it was tempting to label these cells as coccoids, however some of them were observed to divide. To determine their real nature, we used HADA labelling. We overexpressed HdpA from an IPTG-inducible plasmid in strain N6 and did observe coccoid cells and dividing short rods, but also spherical cells with crescent labeling patterns hinting at a different PGN structure (Fig. 4D). This suggests that the overall HdpA level is not the only determinant of coccoid formation, reinforcing the notion that its localization is also an important factor.

### Commitment to the coccoid transition

Since most coccoids were formed between T45-T65 in our conditions, we wondered whether there exists a coccoid commitment timepoint within this period. We tested this by conducting pulse-chase experiments using HADA and RADA to differentiate between cell populations. We grew *H. pylori* cells in the presence of HADA to label the initial population and switched them at either T45 (beginning of transition phase) or T65 (end of transition phase) to fresh medium containing RADA to label the new population. Cells were imaged at T65 (only for T45-switched cells) and T98 (Fig. S8A). First, medium switch at T45, but not at T65, gave rod cells at T98, indicating that fresh medium is able to prevent the transition if provided early in the conversion process. Second, coccoids formed after the T65 medium switch were all exclusively HADA-labeled, indicating that little if any PGN synthesis occurs after T65.

On the contrary, bacterial cells switched at T45 generated at least 4 populations along growth : first (T65), HADA-labeled coccoids (cells that were coccoids or committed at T45), RADA-labeled rods (cells that rapidly grew with extensive PGN remodeling after the medium switch), and RADA/HADA labelled rods (cells that grew slowly or after a lag phase). The latter presumably turned into RADA labeled coccoids only visible at T98. The similar kinetics between the appearance of those RADA labeled coccoids (still a minority 53h after medium exchange) and coccoid appearance in a classical culture suggests that at least part of the cells present at T45 were reset to their initial state by the fresh medium.

Conversely, we grew N6 cells with RADA in exponential phase (T16), switched them to either fresh medium or spent medium obtained from a T65 N6 culture containing HADA. When imaged at T20, HADA-labelled cells in these cultures comprised more coccoids in the case of the switch to spent medium (Fig. S8B).

Altogether, these data indicate that commitment to the coccoid form is a progressive process triggered either by medium exhaustion or the appearance of growth byproducts in which cells slowly lose the ability to grow under the rod form and finally undergo the transition.

### Metabolic adaptation of *H. pylori* through morphological phases

To explore changes in metabolomic profiles of *H. pylori* N6 throughout the morphological transformation, we monitored the metabolome from exponential growth to the late stationary phase using metabolomics approach. Bacteria were grown in liquid cultures from T0 until T165 and captured at five timepoints (T22, T44, T66, T99, and T165) for analysis with LC-MS/MS (liquid chromatography with tandem mass spectrometry) and processing with different metabolomics tools. Multivariate statistical analysis by principal component analysis (PCA) was performed for the five timepoints. The reproducibility and reliability of this experiment was indicated by the PCA score plot showcasing the proximity of each set of biological replicates (except T44_F2) (Fig. S9A).

PCA analysis of the data revealed 3 clusters: T22-T44, T66-T99, and T165, suggesting major metabolomic differences between the pre-transition phase, the post-transition phase, and the late stage . Metabolites that differed significantly between the early (T22 and T44) and late phase (T66, T99, and T165), were determined by partial least squares-discriminant analysis (PLS-DA) (Fig. S9B) and represented as a heatmap (Fig. S9C). Early to late phase transition was characterized by a general downshift in components essential for bacterial growth such as DNA replication (nucleotides and nucleosides), energy production (sugars and amino acids), and membranous structures (phospholipids) (25, 26). On the other hand, we observed an increase in several fatty acids likely indicating elevated activity in certain phospholipid biogenesis pathways.

*H. pylori* metabolites at different time-points were then clustered according to their structure similarity by MS/MS based-molecular networking strategy (27) (Fig. S10). MS/MS datasets were annotated against the GNPS library (28) and the global molecular network was color-tagged according to sample collection time-points. Differences in the level of metabolites by growth phase were consistent with our previous observations. Two clusters containing nucleotides and hexose phosphates were elevated in the exponential phase of growth (Fig. S10A, B). The situation of individual lipids was contrasted, with a drop in phosphatidylglycerol (PG), as observed previously (29), a marginal decrease in phosphatidylethanolamine (PE) but an accumulation of lysophosphatidylethanolamine (LysoPE) in the late phase compared to the early phase (Fig. S10C).

## DISCUSSION

Studies of the *H. pylori* coccoid transition using electron microscopy done on coccoids generated in different conditions concluded either that there are at least two types of coccoids (30, 31) or that coccoids were generated through an intermediate U-shaped form (32–34), without providing definite proof of either hypothesis. Indeed, morphologies of fixed cells observed by electron microscopy could not be assigned with certainty to a step within a morphological pathway. Here, in the particular case of coccoids appearing after prolonged growth, we reconcile the two views by using time-lapse fluorescence microscopy of FDAA labelled cells, which enables us to highlight the PGN and its remodeling within live cells. On one hand we show that indeed two kinds of spherical forms are generated after prolonged growth, stemming from two separate morphological pathways: the coccoid form and the U-form. On the other hand we describe a new intermediate on the coccoid transition pathway. Combining FDAA-, membrane- or DNA-specific fluorescent labelling and cryo-EM, we show that in this intermediate, the lateral PGN is ruptured sufficiently to allow blebbing of the intact inner membrane outside of the PGN sacculus, which mechanically pushes the outer membrane to accommodate the expansion of the cytoplasm. Our data are consistent with the notion that the rupture of the sacculus occurs through hydrolysis of the cross-links between the glycan strands of lateral PGN by the D,D-endopeptidase activity of the HdpA enzyme, the first signs of which are the transversal cracks and the thinning of the sacculus evidenced here by in-liquid AFM. One spectacular feature of the coccoid transition is the fact that the rod shape of the sacculus rapidly morphs into two small (eventually disappearing) protrusions on one side of the spherical bag. This phenomenon could be due either to degradation of the lateral PGN or to a condensation of the sacculus. The intensification of the FDAA signal within these protrusions upon transition and the rod-like shape of the sacculi isolated from coccoids strongly speak in favor of the second hypothesis. What are the forces that drive this PGN “scrunching” The inner membrane-contained cytoplasm escaping from the sacculus tends to adopt an energy-minimizing spherical shape which would expel the sacculus in the absence of other structures. However, the whole system is contained by the outer membrane and the only solution for the sacculus sandwiched between the two membranes is to fold up towards the poles, leading to the characteristic early coccoid shape. This process might be facilitated by the apparent absence of Lpp or β-barrel OM proteins tethering the PGN to the OM (35), consistent with the lack of L,D-transpeptidases in *H. pylori*. In conclusion, our results suggest that coccoids are basically sphaeroplasts in which the PGN is nearly intact except for a slit that enabled its expulsion to a side of the spherical inner membrane.

Interestingly, coccoid transition in *H. pylori* appears similar to the morphological change elicited by β-lactams in *V. cholerae*, including this apparent shrinking of the sacculus, as judged from FDAA-labelling experiments (36). The fact that coccoid transition in *H. pylori* can be triggered by β-lactam treatment, but also simply by prolonged growth suggests that the latter entails a defect in PBP-dependent peptidoglycan synthesis. The imbalance between synthesis and degradation through HdpA activity, made irreversible by subsequent shortening of the peptide stems by Csd4 and Csd6 (7, 8, 24), leads to the rupture in the sacculus.

By contrast, β-lactam treated *E. coli* cells undergo a similar bulging at the beginning but the sacculus does not retract and swelling of the bulge rapidly leads to lysis (23). One possibility to explain the difference with *H. pylori* might be that the *E. coli* PGN does not deform or slide between the membranes as easily because of its anchoring in the OM by Braun’s lipoprotein, which is absent in *H. pylori*. However this does not explain the difference between *E. coli* and *V. cholerae* which also harbours Braun’s lipoprotein.

U-shaped cells have been observed previously and thought to be an intermediate in the coccoid transition. However, the U or doughnut shape of the PGN in a small subset of the spherical cells appearing after prolonged growth suggests that these cells represent the final form of the U-shaped cells. These cells differ from the coccoids as defined here by three features: their appearance does not require the presence of HdpA, they often emit red autofluorescence, suggesting a different metabolical state, and they do not contain DNA, which indicates that they are not viable cells. A possibility is that they represent a DNA source for horizontal gene transfer, an important process in *H. pylori* (37).

*H. pylori* coccoid transition is known to be triggered by different external stresses including starvation (38), as well as by prolonged growth. Whether coccoid transition through prolonged growth depends on external factors (exhaustion of certain nutrients, appearance of growth by-products in the medium), or on an “internal clock” is not known. The fact that cells committed to become coccoids if they were grown without medium change were rescued by the addition of fresh medium at T45 indicates that transition is partly due to external factors. Conversely, the fact that cells in exponential phase partly turn into coccoids without any new PGN synthesis after being switched to spent medium suggest that an endogenous factor is also involved. This factor has to be heterogeneously distributed since in the same experiment, another subset of the cells continues to remodel its PGN. It might also account for the different responses in terms of PGN remodeling observed when switching T45 cultures to fresh medium. Whether it has to do with the state of the PG synthesis machinery, that of the LhiA-HdpA complex (39), the availability of precursors or other factors (24) requires further studies. Actively generated heterogeneity has been shown in other systems to boost the overall fitness of a population (40).

Apart from changes reflecting a low metabolism, the most significant metabolic switch between exponentially growing and coccoid forms pertains to lipid composition: a decrease in phosphatidyl glycerol coupled with an increase in the lysoPE proportion. LysoPE was shown to accumulate upon stress in different bacteria (41–43), to favor membrane curvature, to affect membrane permeability and membrane protein conformation (43, 44). In *H. pylori*, cell subpopulations with a high content of lysoPE, generated through phase variation of the phospholipase A (*pldA*) gene, are more resistant to acidic conditions (45–47). One attractive hypothesis is that coccoid transition constitutes a stress response to which lysoPE participates, possibly changing the interaction of LhiA with the inner membrane with repercussions on HdpA activity, and then stabilizing the membrane bulge after rupture of the sacculus.

In conclusion, whereas some bacterial species adopt a round shape in stationary phase by remodelling their peptidoglycan (48), *H. pylori* coccoid transition follows a different route: the escape of the inner membrane from the sacculus through an endopeptidase-generated crack leads to a sphaeroplast-like cell, with the “empty” peptidoglycan packed on one side of the cell. The preservation of membrane integrity observed here indicates that the load-bearing function of the outer membrane observed in other systems (49) can in certain conditions (probably linked to lipid composition) compensate for the absence of a peptidoglycan shell. The obtained cells are most likely immune to cell wall targeted antibiotics and might therefore be a possible cause of relapse after treatment.

## MATERIALS AND METHODS

### Bacterial Strains and Growth Conditions

*H. pylori* strains N6, 26695, G27, SS1, X47, J99, and B128 were from the lab collection. Bacterial cells were grown on blood agar plates supplemented with 10% defibrinated horse blood and the following antibiotics-antifungal mixture: amphotericin B 2.5 μg⋅mL^−1^, polymyxin B 0.31 μg⋅mL^−1^, trimethoprim 6.25 μg⋅mL^−1^, and vancomycin 12.5 μg⋅mL^−1^. For liquid cultures, we used Brain Heart Infusion (BHI) broth (Oxoid) supplemented with 10% Fetal Calf Serum (Eurobio) and the antibiotics−antifungal mixture without polymyxin B and vancomycin. *H. pylori* cells were grown at 37°C under microaerophilic conditions (6% O_2_, 13% CO_2_, 81% N_2_) using an Anoxomat (MART Microbiology) atmosphere generator and at 180rpm shaking for liquid cultures. From a concentrated glycerol stock stored at - 80°C, *H. pylori* cells were spotted onto a blood agar plate and incubated at the specified conditions for 24h and then transferred to a lawn on a fresh blood agar plate for another 24h in the same condition. Biomass collected from the lawn culture was diluted to OD_600_ of 0.05 to initiate 10 mL liquid cultures in 25-cm^3^ culture flasks (designated as T0). When indicated, 12.5 µM of RADA or HADA (Tocris Bioscience) was added to the liquid medium from T0 until the experiment terminated.

### Growth Kinetics and CFU Assay

Bacterial cultures were initiated according to the standard growth conditions described. At designated timepoints, a 200-µL aliquot was taken from each liquid culture for determination of optical density and for the spot assay. 100 µL of the aliquot was diluted 10-fold in BHI-FCS and measured at 600 nm spectrophotometrically to determine the optical density (OD_600_). The remaining 100 µL was serially diluted 10-fold in BHI-FCS, and 5 µL of each dilution was spotted onto blood agar plates containing 10% FCS, antibiotics-antifungal mixture, and the metabolic dye TTC (2,3,5-Triphenyltetrazolium chloride). Plates were incubated for 5 days, imaged, and CFU counts were recorded.

### Strains

In the *hdpA-*knockout mutant (N6*hp0506*ΩKm), the *HP0506* (*hdpA*) gene is replaced by a non-polar kanamycin resistance cassette (12).

Strains N6 *hdpA-mScartlet-I* and N6 Δ*csd1csd2*-Km were constructed by classical genetic engineering, as detailed in the Supplementary Materials and Methods.

### Microscopy

Fluorescence microscopy was performed with an Axio Observer Z1 inverted microscope (Zeiss) equipped with a Plan Apochromat 100X/1.40 Oil Ph3 M27 Objective of numerical aperture 0.55. Image acquisition was performed using the ORCA-Flash4.0 LT3 digital CMOS camera (Hamamatsu) and ZEN software (ZEN 2.3 pro). Bacterial samples were aliquoted at different timepoints as indicated and were immediately fixed with 70% ice-cold ethanol at -20°C for 15 min, then stored in 20mM HEPES (pH 7) at 4°C until imaging. For imaging of live bacteria, cells were pelleted by centrifugation at 10,000 x g for 1 min and washed once with fresh BHI medium if grown with FDAA. Staining was conducted with 10 µg/mL Hoechst 3342, 10 µg/mL FM4-64fx (Thermo Fisher Scientific), 10 mg/µL WGA-Alexa546, or 10 mg/µL WGA-Alexa488 as indicated. For imaging of autofluorescence signal, non-stained wildtype *H. pylori* N6 was imaged using the mCherry channel (excitation at 555 nm with a 510-560 excitation and 590-4095 emission filter set). Cells were immobilized using 1-mm thick 1% (wt/vol) agarose pads containing physiological sterile water (0.9% NaCl, pH 7) mounted on 1-mm microscopy glass slides and sandwiched in between the agar pad and a glass cover slip.

For time-lapse imaging,*H. pylori* cells were cultured in liquid medium in the presence of RADA as described above until T45. An aliquot was taken and washed once with fresh BHI medium, then diluted to OD_600_ of 0.5 using the same medium. After that, 2 µL of the suspension was deposited on a 35-mm glass bottom µ-dish (Ibidi) and gently spread with the pipette tip. Approximately 1-mm thick of a 1.5% agarose-BHI pad (BHI supplemented with 10% FCS and 1.5% ultrapure agarose (Invitrogen, Cat #16520050)) was placed on top to immobilize the cells. The µ-dish was placed at 37°C, under a 10% CO2, 5% O2, 90% humidity atmosphere in an environmental chamber (PECON Incubator PM S1 – TempModule S, CO2 Module S, O2 Module S, and Heating Insert P-Labtek S1) . Image acquisitions were conducted every 30 min for a period between 8-24 hours for phase contrast and RADA staining (excitation with a 555-nm laser with a 533-558 and 570-640 filter set). Processing of the images was done using ImageJ.

### Image Analysis with ImageJ

Image processing and analyses of fluorescence intensity profiles were conducted using FIJI (version 2.9.0) and the MicrobeJ plugin (version 5.13). Size measurements of the 8 *H. pylori* strains were conducted on phase contrast images. Cells were detected using the following parameters: 1) Rod cells were detected with a fit-shape rod-shaped model of a length between 1-5 µm and width between 0.1-1 µm; and 2) Coccoid cells were detected with a fit-shape circle model of a length and a width between 0.5-1.5 µm. The resulted measurements were exported and visualized using GraphPad. For demographic analysis of fluorescence intensity profiles, bacteria were detected using contour or medial axis dilated by 0.1 µm. Clusters of 2 and more cells were excluded from the analysis and only isolated bacteria were included. Each analysis was conducted using triplicates.

### Image Analysis with ilastik

Pipelines for morphological analysis of bacteria were established using the pixel classification and object classification [inputs: raw data, pixel prediction map] workflows in ilastik (version 1.4.0). Raw images were flattened when necessary and converted to tif files in FIJI before uploading to ilastik. Images were divided into a training set, test set, and data set. The training set was used to establish and train a pipeline whose performance was assessed using the test set. Once validated, the trained pipeline was use to analyze images in the data set for generating results. Phase contrast images (single channel) were used to establish the phase designation pipeline, RADA images (single channel) were used to establish the morphological analysis pipeline, HADA images (single channel) were used for quantification of the U-form, and images of HADA with autofluorescence (two channels flattened) were used for quantification of autofluorescent cells.

Signal (phase or fluorescence) was distinguished from background using the pixel prediction workflow through reinforcement learning to eventually generate pixel probability maps. These were then used to conduct object classification along with their corresponding tif files. The threshold and filter were set to 50 minimum and 10000 maximum using Hysteresis method, and all object feature selections were included. Depending on the pipeline, between 3-4 morphological classes were included as indicated and the software was trained to distinguish between each class through reinforcement learning. When a satisfactory result was obtained, object identities and feature tables were exported.

Once the pipeline had been established, its performance was then assessed using the test set. Tif files, pixel probability maps, object identities, and feature tables were generated for all images in this set using the trained pipeline. A confusion matrix was constructed based on the generated predicted classes and separately assigned labels on the same cells. The pipeline was retrained if its calculated accuracy was below 90% or used to analyze the data set if it was determined to be above. Depending on the analysis, at least three replicates or three separate experiments of data sets were included.

### Pulse-Labeling with FDAA

Generally, bacteria were grown with or without FDAA (12.5 µM) then captured at various timepoints for labeling or exchanging FDAA. For short pulse-labeling in the presence of antibiotics (meropenem or aztreonam), *H. pylori* cells were cultured with either RADA or HADA (12.5 µM) from T0. Then, they were captured at T20 for removal of the probe by washing twice with BHI medium followed by resuspension in the same volume of fresh BHI medium. Multiple 500 µL aliquots were taken from the suspension and transferred into a 24-well plate along with 1X, 2X, or 4X of MIC of one of the antibiotics except for controls (untreated). The plate was then incubated for 3 hours in the standard condition. Next, 100 µM of the counterpart probe (HADA if RADA at T0 and vice versa) was added to cultures and they were allowed to grow for another 20 minutes, fixed with 70% EtOH and stored for imaging.

For pulse-labeling with with medium exchange, bacteria were grown with HADA (12.5 µM) from T0. When captured at T45 or T65, the entire culture was collected after a small aliquot was taken for fixation, washed twice with fresh BHI medium, then resuspended in the same volume of fresh BHI medium before splitting into two separate cultures. One was given fresh RADA (12.5 µM) while the other was kept as control. After that, the bacterial cells were allowed to grow again until T65 (if captured at T45) and T90 followed by ethanol fixation and imaging.

For the experiments with spent medium, RADA-labeled cells (or non-labeled controls) were collected from T16 cultures. They were resuspended either in fresh medium (i.e. medium from a T16 culture in HADA) or in spent medium (medium from a T65 culture in HADA), supplemented with fresh HADA (12.5 uM). These cells were then transferred into 24-well plates, grown for 4 hours and fixed with 70% ethanol for imaging.

### Cryo-Electron Microscopy and Tomography

Sample preparation: The protocol for grid preparation was adapted from previously described workflows (50, 51). Glutaraldehyde-fixed cell suspension of *H. pylori* was supplemented with 10 nm protein-A gold immediately before application to freshly glow discharged Quantifoil R3.5/1 gold 200 mesh grids and was plunge-frozen with a Vitrobot Mark IV.

Data collection: Two-dimensional images of cells and tilt series were collected on a Titan Krios transmission electron microscope operating at 300 kV fitted with a K3 direct electron detector (BioQuantum energy filter (slit width 20 eV)). Collection of cellular images was performed at a pixel size of 15.65 Å. Tilt series were collected at a pixel size of 3.37 Å with defoci ranging from -6 to -12 μm, tilts ranging from -60° to +60° with a tilt increment of 1° and a final total dose of 140 e^−^ Å^−2^.

Data analysis: Tomogram were reconstructed using Relion-5 (52) Alignment of tilt series was performed using IMOD (53) by automatic fiducial based alignment for rod cells and AreTomo (54) for coccoid cells. Final image data were analysed using Fiji (55).

### Sacculi Extraction for Fluorescence Imaging

Liquid bacterial cultures were grown until T20 (for rods) or T90 (for coccoids) in the presence of RADA and a total of 20-unit equivalent of OD_600_ was collected via centrifugation at 20,000 x g for 15 min followed by resuspension in 1 mL of 0.25% SDS. The suspension was boiled for 20 min at 100°C and then centrifuged at 20,000 x g for 5 min. The resulting pellet was washed 3 times with 1.5 mL distilled water via centrifugation, and finally resuspended in 1 mL of distilled water. After that, 0.5 mL of TRIS/HCl buffer (0.1 M, pH 6.8) containing 15 µg/mL DNase and 60 µg/mL RNase was added, and the mixture was incubated for 60 min at 37°C with 800 rpm agitation. Subsequently, 0.5 mL of TRIS/HCl buffer (0.1 M, pH 6.8) containing 50 µg/mL trypsin added and the mixture was incubated for another 60 min in the same condition. Then, the suspension was boiled for 3 min at 100°C in a heating block followed by centrifugation at 20,000 x g for 5 min. The pellet was washed once with 1 mL distilled water via centrifugation and then resuspended in 20 µL of distilled water and stored at 4°C.

### Sacculi Extraction for AFM

Liquid bacterial cultures were grown until T20 (for rods) or T90 (for coccoids) and a total of 500-unit equivalent of OD_600_ was collected via centrifugation at 15,950 x g for 10 min followed by resuspension in 10 mL of fresh BHI. The suspension was added dropwise into 10 mL of 5% (w/v) SDS preheated to 100°C and kept at 100°C for 30 min. The pellet was collected from the suspension by ultracentrifugation at 400,000 x g for 15 min at room temperature and washed 4 times with H_2_O via ultracentrifugation. Next, the pellet was resuspended in 3.6 mL sodium phosphate buffer (50 mM, pH 7.3) and 0.4 mL of trypsin (1 mg/mL) was added. The mixture was incubated at 37°C overnight with 800 rpm agitation. Then, SDS was added to the mixture to a final concentration of 2% and it was boiled at 95°C for 30 min. Once the suspension cooled down to room temperature, it was centrifuged at 13,000 rpm for 5 min and its supernatant was ultracentrifuged at 400,000 x g for 30 min. Subsequently, the resulting pellet was washed twice with distilled water via ultracentrifugation before finally resuspended in 100 µL of ultrapure water and stored at 4°C.

### Atomic Force Microscopy imaging of purified *H. pylori* PGN

All high-resolution AFM images were captured in liquid (imaging buffer: 10 mM Tris, pH:7.8) using Fastscan-D cantilevers in Amplitude Modulation Intermittent Contact Mode (also known as Tapping mode) on a Dimension FastScan Bio (Bruker, Santa Barbara). The imaging parameters are as follows; Drive frequency: 70 - 100 kHz; Scan rate: 1 Hz; Scan angle: 0°; and Amplitude setpoint: 2.5 – 3.5 nm (70 – 80 % of the free amplitude). Image processing and analysis used both mean plane subtraction and measure distance function in Gwyddion 2.56 software (56). Immobilization of *H. pylori* PGN for AFM and image processing are described in the supplementary Materials and Methods. For thickness measurements, AFM height topographic images of dehydrated PGN were captured in air using AFM tapping mode with Nunano SCOUT 350 - Silicon AFM probe (spring constant: 42 N/m, Resonance frequency: 350 kHz) at a free amplitude of 10 nm on a Dimension FastScan Bio (Bruker, Santa Barbara).

### Culturing of *H. pylori* N6 for LC-MS/MS

Cultures of *H. pylori* were started as above. The bacteria from agar plate were precultured in BHI + 10% FCS under microaerobic condition at 37°C, 180 rpm for 22h and an aliquot was withdrawn and diluted to an optical density at 600 nm (OD_600_) of 0.05 in a fresh culture medium. A 1 mL sample was withdrawn at T22, T44, T66, T99, and T165. The amounts of bacteria were normalized to an OD_600_ of 2.15 ± 0.15. Each experiment was repeated in five replicates.

### Intracellular metabolites extraction

Bacteria from the 1 mL samples were harvested by centrifugation at 16000 x g for 5 min at 4°Cs. The pellet was washed with HEPES 1 mM solution and centrifuged again for 5 min at 16000 x g at 4°C. After removing the supernatant, 1 mL of cold methanol/ethanol/water (5/3/2, −20°C) was added to the pellet. The tubes were then immersed in liquid nitrogen for metabolism quenching and incubated at 4°C for three hours. The tubes containing bacteria in the extraction solution were vortexed and then centrifuged 15 min at 16000 x g at 4°C. A 900 μL aliquot of the supernatant was collected and vacuum-dried using a SpeedVac instrument (Jouan RC1010) and stored at −80°C until analysis.

### Metabolites LC-MS/MS profiling

Dried samples were dissolved in 200 μL of 10 mM ammonium carbonate buffer (pH 10, adjusted with ammonium hydroxide)/acetonitrile (20/80, v/v). Tryptophane-d_5_ (3 µg/mL, Sigma-Aldrich) was added to monitor the LC–MS performance. A quality control (QC) sample was prepared by pooling all extracts from different time points sampling. This QC sample was injected regularly throughout the acquisition batch.

LC-ESI-MS/MS analysis was performed by UHPLC-HRMS, using a Dionex Ultimate 3000 UHPLC coupled to a qExactive Focus mass spectrometer (Thermo Fisher Scientific) with a capillary voltage at −3 kV in the negative ionization mode and a capillary temperature set at 280 °C. The sheath gas pressure and the auxiliary gas pressure (nitrogen) were set at 60 and 10 arbitrary units, respectively. The detection was performed from *m/z* 80 to 1200 using a resolution set at 70 000 at m/z 200 (full width at half-maximum), with an automatic gain control (AGC) target of 3 × 10^6^ ions and an automated maximum ion injection time (IT). Data-dependent MS/MS were acquired on a “Top 3” data-dependent mode using the following parameters: resolution 35000; AGC 1×10^5^ ions, maximum IT 50 ms, stepped NCE of 10%, 15% and 30%. The mass spectrometer was calibrated externally before each analysis in both ESI polarities using the manufacturer’s predefined methods and the recommended calibration mixture provided by the manufacturer.

The chromatographic separations were performed using a Sequant ZIC-pHILIC (5 μm, 2.1 × 150 mm) column at 20 °C (Merck, Darmstadt, Germany) operated under gradient elution at 0.2 mL/min. The mobile phases used were water with 10 mM ammonium carbonate, pH 10, adjusted with ammonium hydroxide (A) and acetonitrile (B). Elution started with an isocratic step of 2 min at 80% B, followed by a linear gradient from 80 to 40% of phase B from 2 to 12 min. The chromatographic system was then rinsed for 5 min at 0% B, and the run was ended with an equilibration step of 15 min (57). The autosampler compartment temperature was set at 4°C and the injection volume was 5 µL.

### Data analysis

The raw data files (Thermo Fisher.raw) were converted to standard data-format to mzML format using the MSConvert software, part of the ProteoWizard package (58). All mzML files were then processed using MZmine 4.0.3 (59) and the online Workflow4Metabolomics.org platform (60). Detailed parameters of data processing are available in the Supplementary Materials and Methods.

## ACKNOWLEDGMENTS

The Boneca laboratory was supported by the following programs: Laboratoire d’Excellence “Integrative Biology of Emerging Infectious Diseases” (ANR-10-LABX-62-IBEID) and Équipe FRM Grant (EQU202403018034).This work was also funded by the Medical Research Council, as part of United Kingdom Research and Innovation [Programme MC_UP_1201/31 (T.A.M.B)], by the Wellcome Trust (grants 225317/Z/22/Z (T.A.M.B), 212197/Z/19/Z (SJF, JKH), 104110/Z/14/A (SJF, JKH)) and by the UK Research and Innovation (UKRI) Strategic Priorities Fund grant EP/T002778/1 (JKH, SJF). T.A.M.B. and O.E.R.S. would like to thank the Lister Institute for Preventative Medicine for support and the MRC LMB electron microscopy and scientific computing facilities. For the purpose of open access, a CC-BY public copyright licence has been applied by the authors to the present document and will be applied to all subsequent versions up to the Author Accepted Manuscript arising from this submission.

