## Supplemental Information for "Morphological transformation in *Helicobacter pylori* is a dynamic process leading to two types of coccoid"

**This PDF file includes:**

Supporting Information text
Figures S1 to S10
Tables S1 to S2
SI References

**Other supporting materials for this manuscript include the following:**

Movies S1 to S4

### Supporting Information Text

### Material and Methods

#### Bacterial Strains and Growth Conditions

*H. pylori* strains N6, 26695, G27, SS1, X47, J99, and B128 were from the lab collection. Bacterial cells were grown on blood agar plates supplemented with 10% defibrinated horse blood and the following antibiotics-antifungal mixture: amphotericin B 2.5 μg·mL−1, polymyxin B 0.31 μg·mL−1, trimethoprim 6.25 μg·mL−1, and vancomycin 12.5 μg·mL−1. For liquid cultures, we used Brain Heart Infusion (BHI) broth (Oxoid) supplemented with 10% Fetal Calf Serum (Eurobio) and the antibiotics−antifungal mixture without polymyxin B and vancomycin. *H. pylori* cells were grown at 37C under microaerophilic conditions (6% O2, 10% CO2, 84% N2) using an Anoxomat (MART Microbiology) atmosphere generator and at 180rpm shaking for liquid cultures. From a concentrated glycerol stock stored at -80°oC, *H. pylori* cells were spotted onto a blood agar plate and incubated at the specified conditions for 24h and then transferred to a lawn on a fresh blood agar plate for another 24h in the same condition. Biomass collected from the lawn culture was diluted to OD600 of 0.05 to initiate 10 mL liquid cultures in 25-cm3 culture flasks (designated as T0). When indicated, 12.5 µM of RADA or HADA (Tocris Bioscience) was added to the liquid medium from T0 until the experiment terminated.

#### Growth Kinetics and CFU Assay

Bacterial cultures were initiated according to the standard growth conditions described. At designated timepoints, a 200 µL aliquot was taken from each liquid culture for determination of optical density and for the spot assay. 100 µL of the aliquot was diluted 10-fold in BHI-FCS and measured at 600nm spectrophotometrically to determine the optical density (OD600). The remaining 100 µL was serially diluted 10-fold in BHI-FCS, and 5 µL of each dilution was spotted onto blood agar plates containing 10% FCS, antibiotics-antifungal mixture, and the metabolic dye TTC (2,3,5-Triphenyltetrazolium chloride). Plates were incubated for 5 days, imaged, and CFU counts were recorded.

#### Strains

The N6 *hdpA*-*mScartlet-i* strain was obtained by natural transformation of the N6 strain with a DNA fragment containing a fusion of the 3' end of the hdpA gene with the *m-Scarlet-i* gene followed by a kanamycin resistance cassette and the 5' end of the *HP0507* gene, enabling homologous recombination and selection.

In wildtype N6 (and 26695), *hdpA* and the following gene, *HP0507*, are most likely translationally coupled through a TAATG sequence (where the undescored and bold bases correspond to the STOP codon of the first gene and the initiator codon of the second, respectively). We have preserved this organization in our construct.

First, plasmid pUC18 *mSc*-K2 harboring a translationally coupled *m-Scarlet-i-aphA3* operon was obtained by co-transformation in DH5a of the products of 2 PCR: that of plasmid pTriEx-RhoA-wt_mScarlet-i_SGFP2 (1) gift from Dorus Gadella (Addgene plasmid #85071; http://n2t.net/addgene:85071 ; RRID:Addgene_85071)) using primers Kan-mSc and mSc-Kan, and that of pUC18-K2 (2) using primers KanG2 and KanD2. Then, to generate a F('*hdpA-mScarlet-i*) *aphA3 HP0507*' fragment, pUC18 mSc-K2 was subjected to a PCR using primers 506-mSc and mSc-507 while flanking regions were generated by PCR of N6 genomic DNA with primers 506F1 and 506R1 for the '*hdpA* side and 507F1 and 507R1 for the HP0507' side; the 3 amplicons were combined in a final PCR using primers 506F1 and 507R1 to generate the final fragment. In the final construct: *hdpA* and *mScarlet-i* are fused in one gene encoding a protein in which the C-terminal residue of HdpA is followed by the initiator methionine of *mScarlet-i*; *mScarlet-i* and *aphA3* are translationally coupled through an ATGA sequence while the *aphA3 HP0507* junction consists of the same TAATG sequence as the *hdpA HP0507* junction in the wt genome.

The double knockout mutant of *csd1-csd2* genes (N6Δ*csd1csd2*-Km) was constructed through natural transformation of PCR products consisted of a kanamycin resistant cassette flanked by the homology region upstream of *csd2* (*folC*) and downstream of *csd1* (*ccmA*). The upstream and downstream homology regions were amplified from the genome of strain 26695 using primer folC_F (5'-GATTATATTGAACGCTACC-3') and overlapping primer folC-Kana_R (5'-ACCCGGGTACCTCAATCCCTCTTCTC-3') and overlapping primer Kana-ccmA_F (5'-ACCCGGGGATCCGCAATCTTTGATAACAATAATAAATCG-3') and primer ccmA_R (5'-ATCCTTACAAGCGAGTATTTGG-3'), respectively. The kanamycin resistant cassette was amplified from pUC18 with primers Puc18_Kana_R (5'-GGATCCCCGGGTCATTATTCC-3') and Puc18_Kana_F (5'-GGTACCCGGGTGACTAACTAGG-3') and fused with the flanking homologies of *csd1-csd2* via overlap PCR using primers folC_F/ccmA_R.

**Cryo-Electron Microscopy and Tomography**

Sample preparation: The protocol for grid preparation was adapted from previously described workflows (3, 4). Briefly, 1 µL of 10 nm protein-A gold (CMC Utrecht) was added to 12 µL of the 2.5% glutaraldehyde-fixed cell suspension immediately before grid preparation. 2.5 µL of this mixture was applied to freshly glow discharged Quantifoil R3.5/1 gold 200 mesh grids and plunge-frozen using a Vitrobot Mark IV (ThermoFisher) following a wait time of 2 s with a blot force of -10 and blot time of 6 s. Grids were stored in liquid nitrogen until further investigation.

Data collection: two-dimensional images of cells as well as tilt series were collected on a Titan Krios transmission electron microscope (ThermoFisher) operating at 300 kV fitted with a K3 direct electron detector (Gatan) and BioQuantum energy filter (Gatan, slit width 20 eV). Collection of two-dimensional cell images was performed with the EPU software at a nominal magnification of 11,500 x (resulting in a pixel size of 15.65 Å). Tilt series collection was performed with the SerialEM software at a magnification of 26,000 x (resulting in a pixel size of 3.37 Å) with a defocus range of -6 to -12 µm between -60° to +60° with a tilt increment of 1°.

Data analysis: Tilt series were reconstructed into tomograms using the Relion-5 pipeline (5). Alignment of tilt series was performed using IMOD (6) by automatic fiducial based alignment for rod cells and AreTomo (7) for coccoid cells. Tomograms and two-dimensional images were analysed using Fiji (8). A bandpass filter was applied to all images.

**Immobilization of purified *H. pylori* PGN and AFM imaging**

Before AFM imaging, purified PGN was immobilized on a poly-l-ornithine-coated mica substrate. Briefly, freshly cleaved mica was incubated with 40 µl of poly-l-ornithine (5 µg/ml in water) for 5 min. Next, the substrate was rinsed three times with HPLC-grade water and dried with nitrogen flow. Afterward, 40 µl of purified PGN was added onto poly-l-ornithine-coated mica substrate and incubated for 10 min and then rinsed three times with HPLC grade water and dried with nitrogen flow.

**AFM image processing and analysis**

All high-resolution images were leveled by mean plane subtraction followed by line flattening using Gwyddion 2.56 software. To measure the thickness of the dehydrated PGN, we employed thickness measurement method described in our previous work (9). For the cell width measurement, we utilized the "measure distances and directions between points" tool in Gwyddion to measure the short axis of the cell.

**Metabolomics data analysis**

*MZmine LC-MS/MS Processing and Molecular Networking Parameters*

The mass detection was realized using factor of lowest signal algorithm with noise factor as 5 for MS1 and 2.5 for MS2. The chromatogram builder was used using a minimum group size of scans of 4, a minimum intensity for consecutive scans of 2.0E3, a minimum highest intensity of 1.0E4 and m/z tolerance of 0.002 m/z or 10 ppm. The chromatogram deconvolution was performed using the local minimum feature resolver with the following settings: chromatographic threshold = 90%, minimum search range RT/Mobility = 0.05, minimum absolute height = 1.0E4, minimum ratio peak top/edge = 5, peak duration range = 0 – 1.2 min. Isotopes were grouped using the isotopic peaks grouper algorithm with an m/z tolerance of 0.0015 or 3 ppm and an RT tolerance of 0.02 min. The peak alignment algorithm was used with the following settings: m/z tolerance of 0.0015 or 5 ppm, weight for m/z of 3, retention time tolerance of 0.1, and weight for RT of 1. The resulting peak list was filtered to keep only rows with MS2 features and feature is kept if it was detected in more than 3 samples. The .mgf and .csv (for RT, m/z, peak areas) files were exported using the dedicated “Export/Submit to GNPS/FBMN” option. A network was then created using GNPS2 platform where edges were filtered to have a cosine score above 0.7 and more than six matched peaks. Further edges between two nodes were kept in the network if and only each of the nodes appeared in each other′s respective top ten most similar nodes. The spectra in the network were then searched against GNPS spectral libraries. All matches kept between network spectra and library spectra were required to have a score above 0.7 and at least six matched peaks. The molecular networking data were analyzed and visualized using Cytoscape v.3.10.220. The GNPS job is accessible at: https://gnps2.org/status?task=9f5100b61708450c9f8d2ea8a249670c

*Data analysis using Workflow4Metabolomics.org platform*

Data pre-processing: Peak detection and integration were performed using the CentWave algorithm of the XCMS package (10) integrated on the Workflow4Metabolomics.org platform (11). Annotation of isotope peaks, adducts and fragments was realized by CAMERA annotation (12).

Normalization: signal drift was corrected by local polynomial regression models (LOESS) to quality control samples (13). The features generated from XCMS were retained only if they met the following criteria: (i) the feature coefficient of variation in QC samples is less than 30% and (ii) the peak area mean ratio of blank samples to biological samples is less than 0.1.

*Metabolite Annotation*

Annotation was performed by using MoNA database including Massbank, HMDB, LipidBlast and GNPS with a masse tolerance of 5 ppm of the precursor and 10 ppm of the product.

*Statistical analysis*

Multivariate data analyses such as principal component analysis (PCA) and partial least-squares discriminant analysis (PLS-DA) were performed to highlight discriminant metabolites implicated in different bacterial growth phases. To ensure the robustness of the prediction, variables were gradually excluded according to results obtained from Variable Importance in the Projection (VIP) plot and univariate p-values (non-parametric Wilcoxon-Mann-Whitney statistical test with a Benjamini-Hochberg false discovery rate (FDR) correction)). The metabolites were retained only if their VIP was greater than 1.29 and the p-value was less than 0.05. All discriminant metabolites with annotations were uploaded to MetaboAnalyst (http://www.metaboanalyst.ca/) to generate a heatmap.

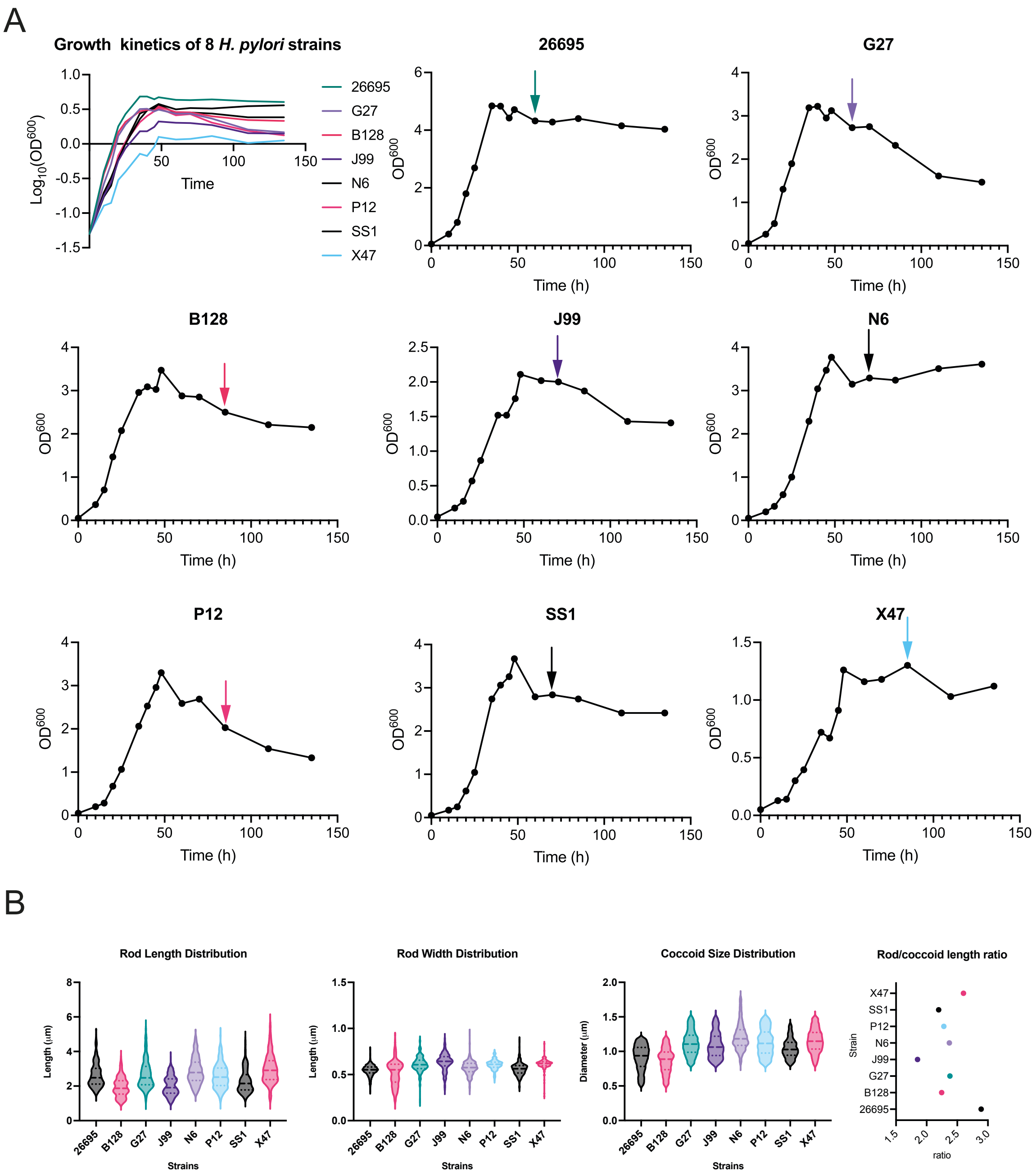


**Fig. S1.** Comparison of 8 strains of *H. pylori*. A) Growth curves of all 8 strains on a log scale and individual growth curves of each strain with their coccoid formation initiation timepoint indicated by an arrow. B) Size distributions of the helical-rod and coccoid form. Distributions of rod length and rod width were measured from phase contrast images of bacterial cells at T10, and distribution of coccoid diameter was measured from phase contrast images of bacterial cells at T135. Rod to-coccoid length ratio was calculated using the mean rod length and the mean diameter of the coccoid for each strain.

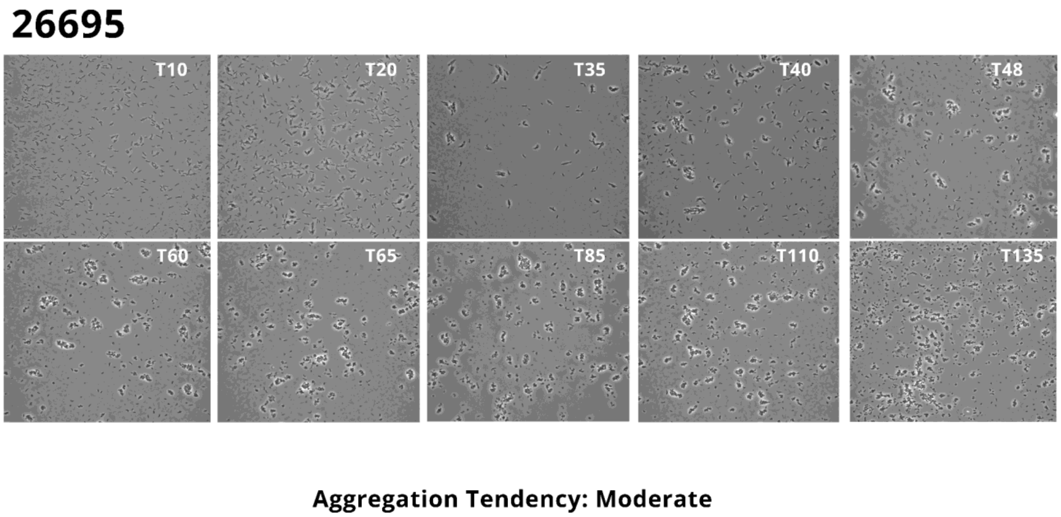


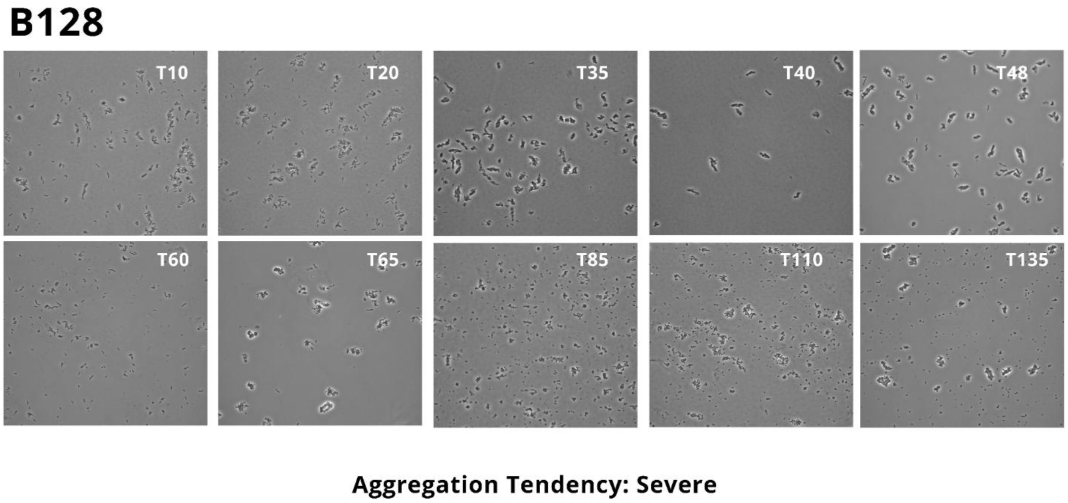


**Fig. S2.** Comparison of 8 strains of *H. pylori*. Representative microscopy images of liquid cultures in standard conditions for the 8 tested strains at indicated time-points..

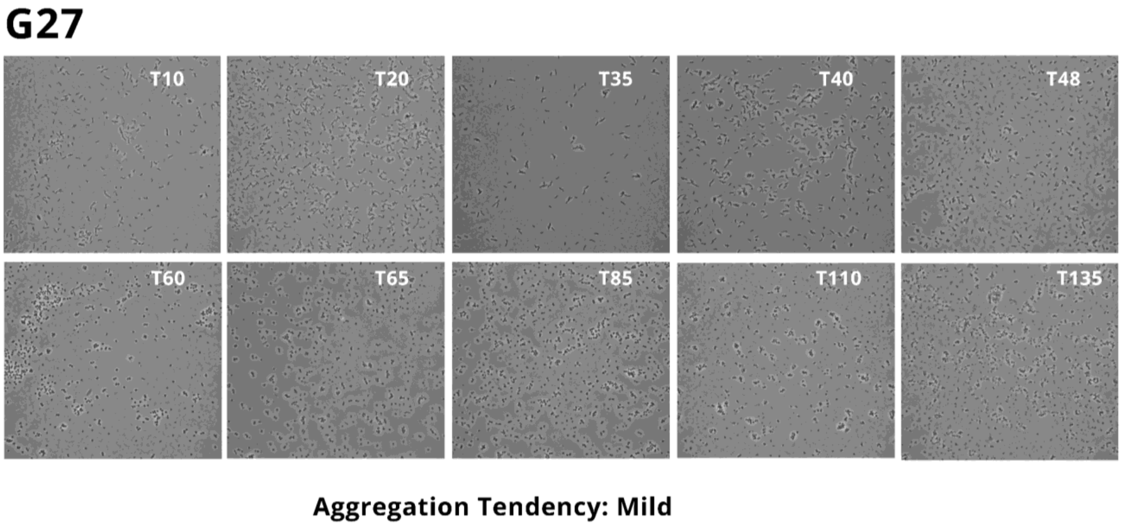


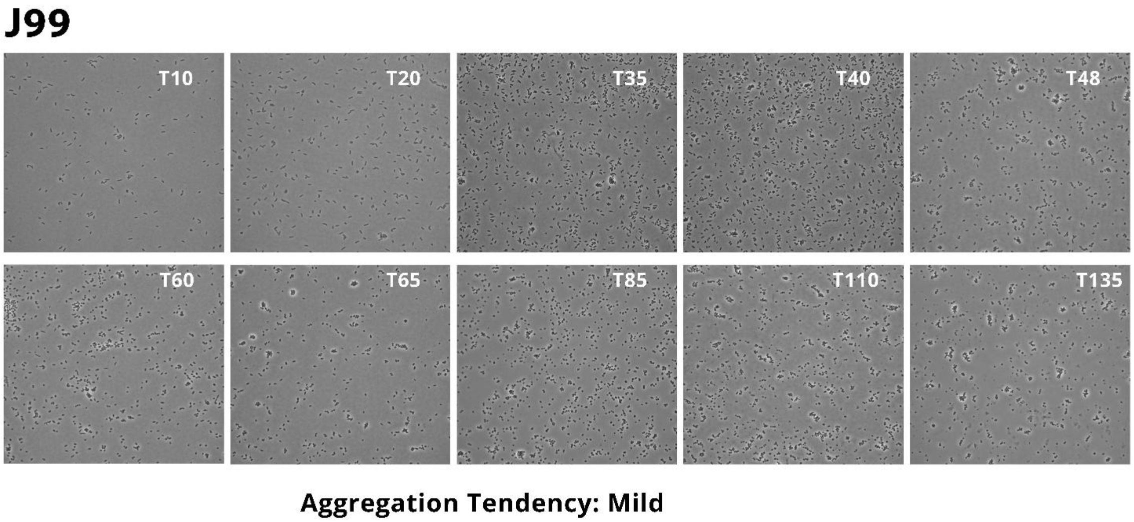


**Fig. S2.** Continued.

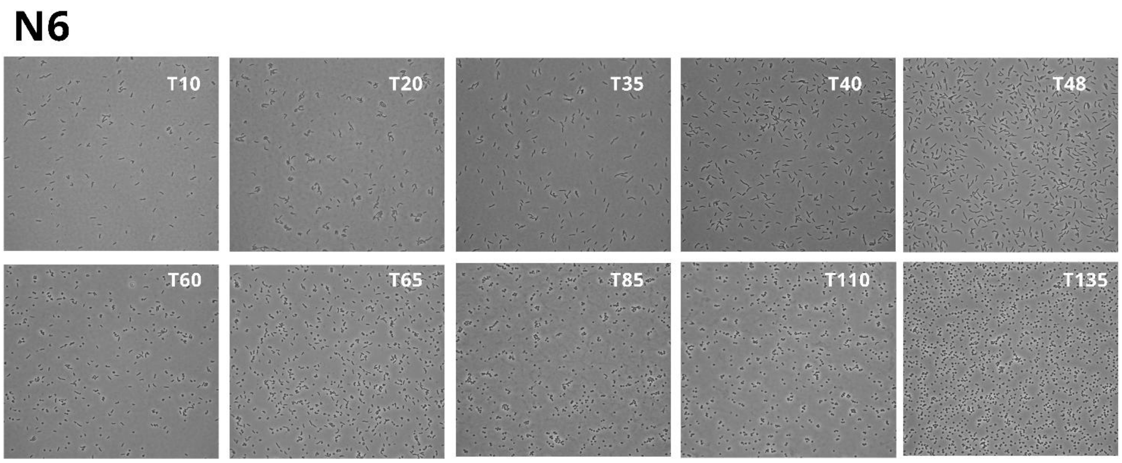


Aggregation Tendency: Mild

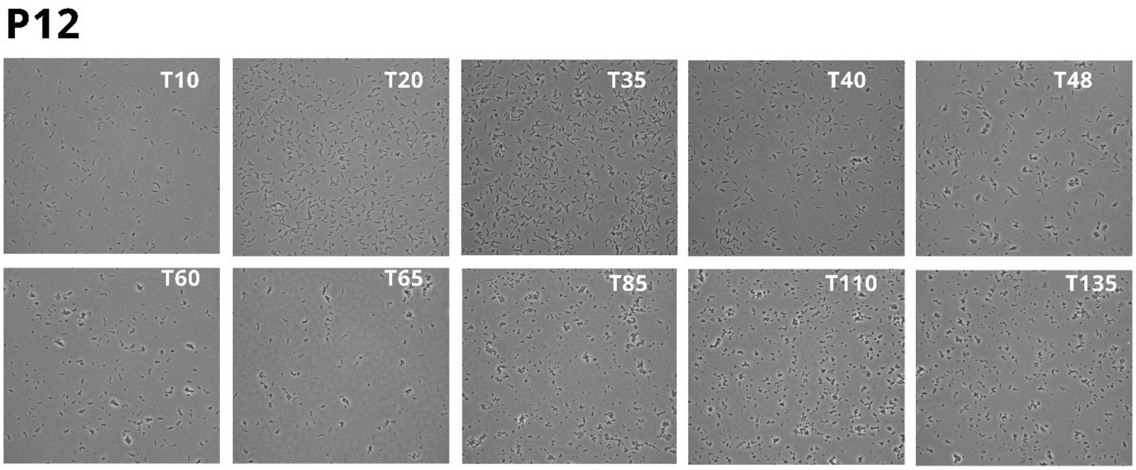


Aggregation Tendency: Moderate

**Fig. S2.** Continued.

## SS1

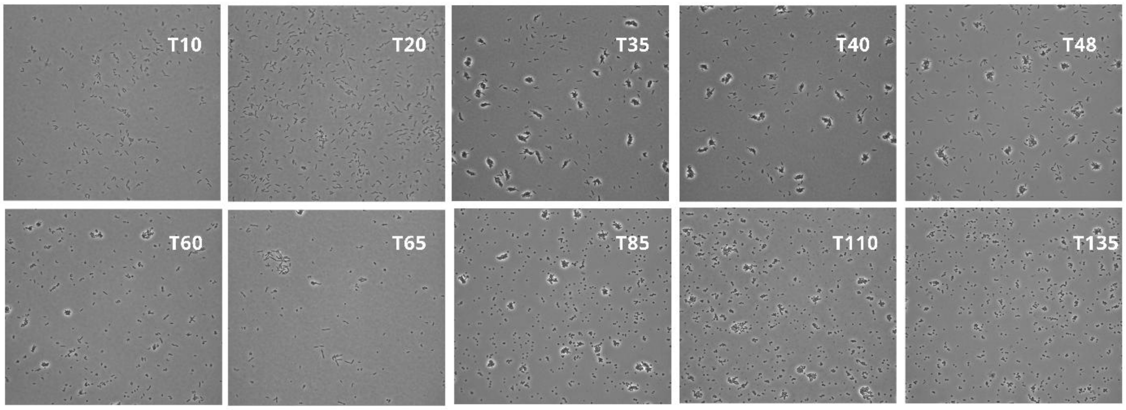


**Aggregation Tendency: Moderate**

## X47

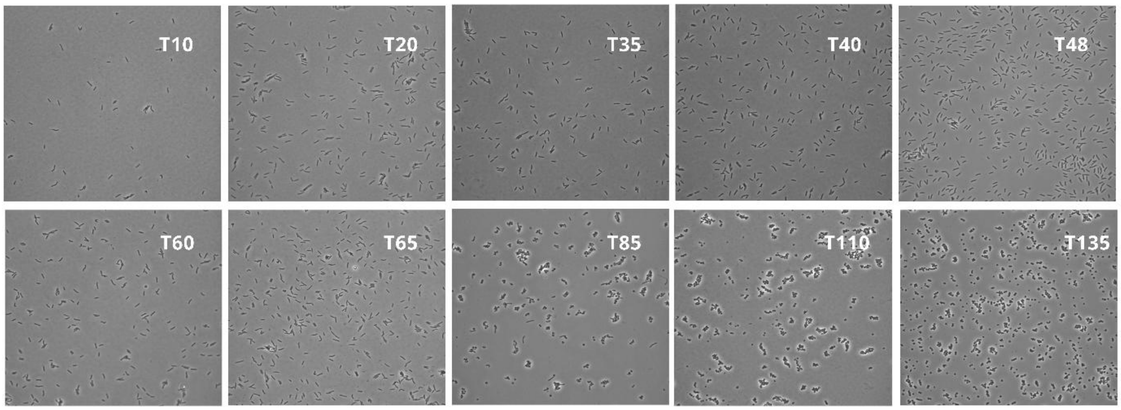


**Aggregation Tendency: Moderate**

**Fig. S2.** Continued

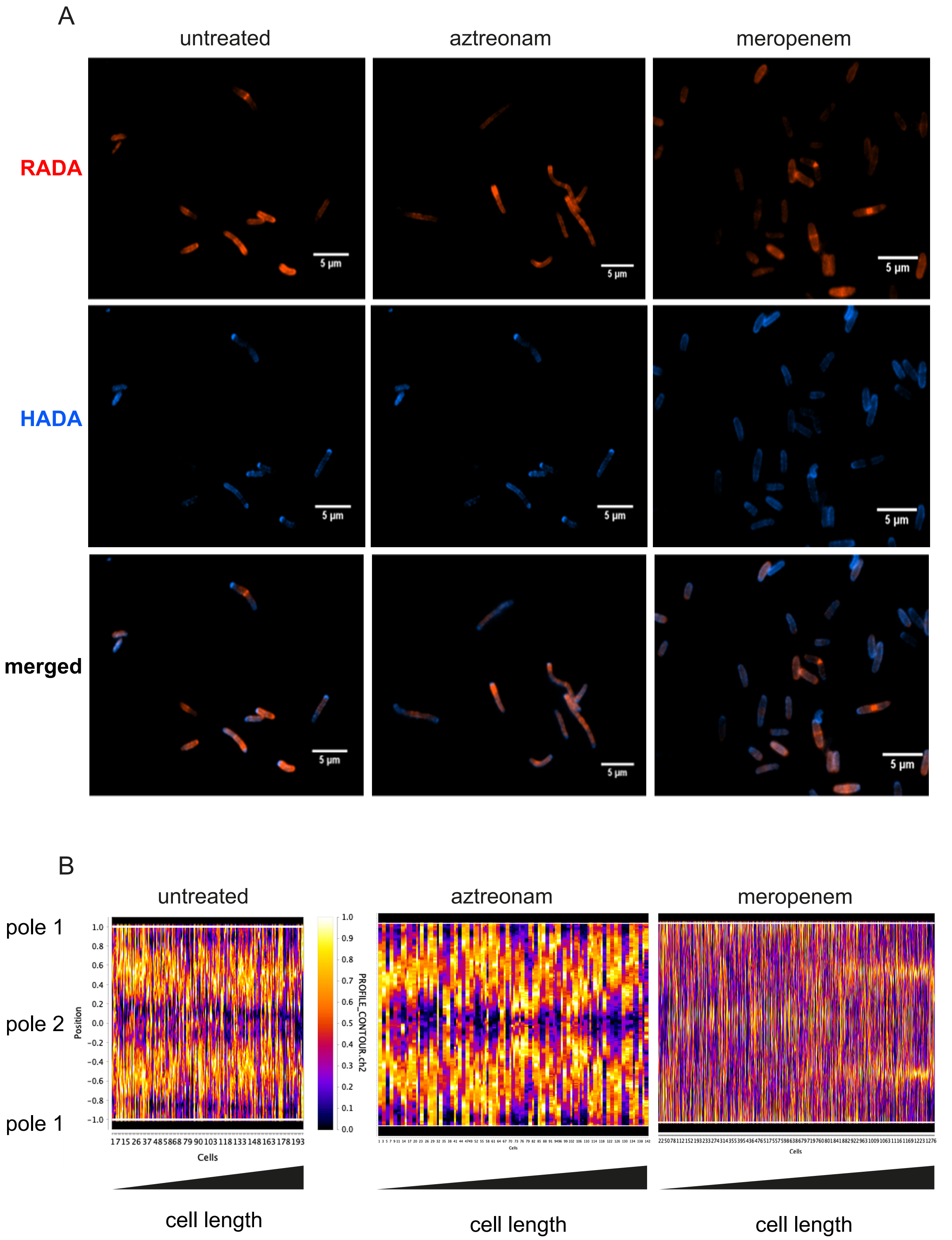


**Fig. S3.** FDAAs incorporation mechanism in *H. pylori*. A) Effects of beta lactams on morphology and FDAAs incorporation in *H. pylori*. Fluorescence images of *H. pylori* N6 cells stained with HADA (blue), treated with the indicated antibiotic and chased with RADA (red). B) Demographs of RADA fluorescence intensity along cell contour, from designated pole 1 to pole 2 and back to pole 1, sorted by cell length.

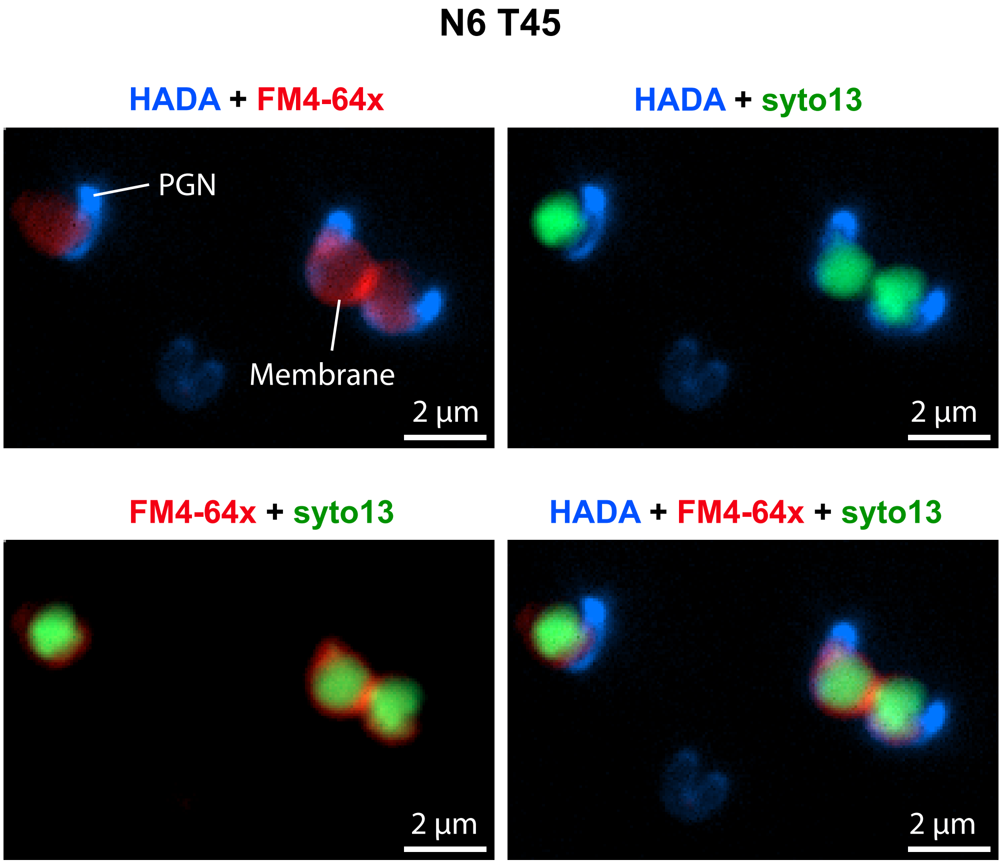


**Fig. S4.** Overlay fluorescence images of *H. pylori* cells at T45 grown the presence of HADA (blue) and stained with FM4-64x (red) and Syto-13 (green). .

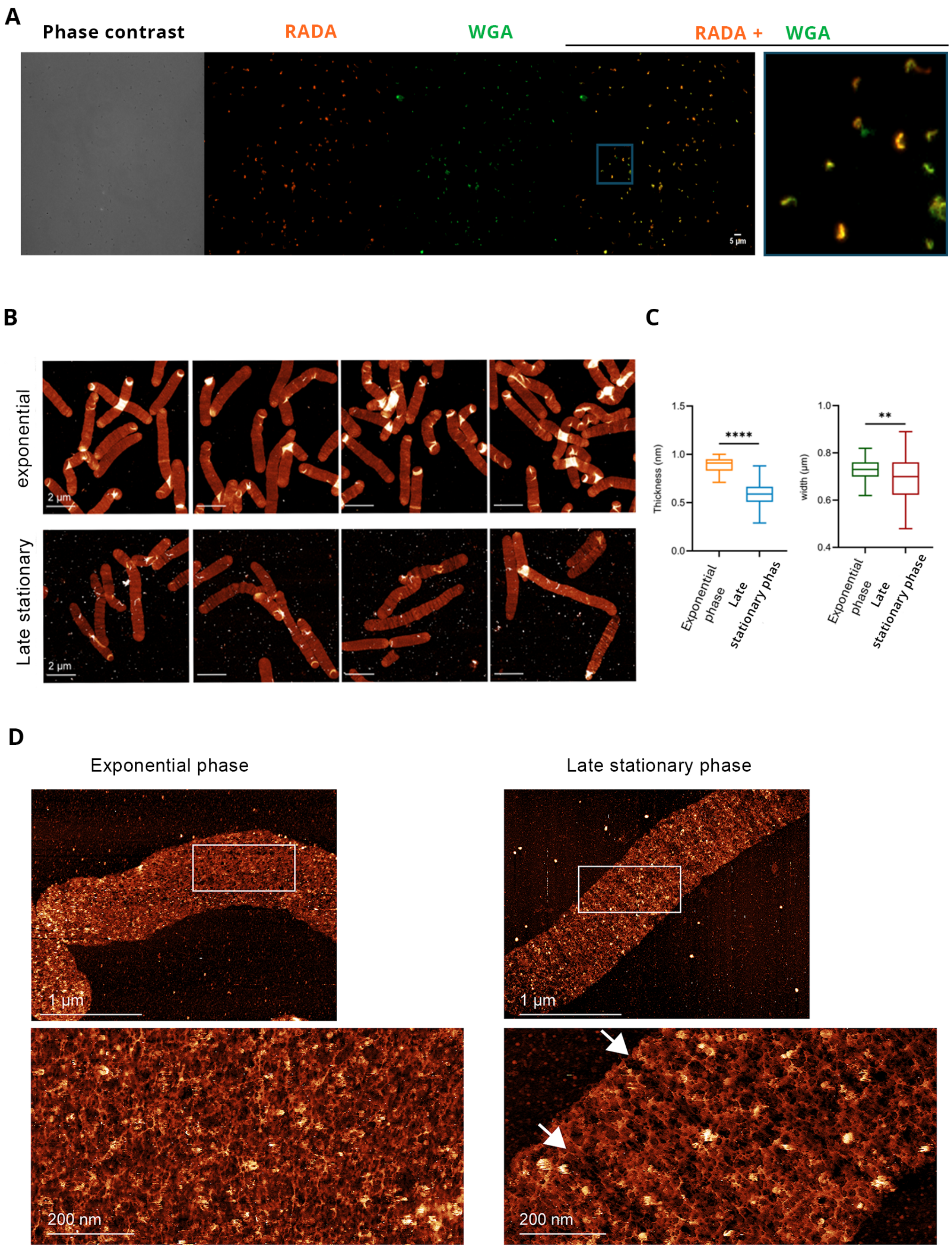


**Fig. S5.** Sacculi of *H. pylori* rod and coccoid forms. A) sacculi of *H. pylori* cells grown in the presence of RADA (red) until T90 and stained with WGA (green). B) low-resolution AFM micrographs collected in air of *H. pylori* sacculi at exponential (T20) and late stationary phase (T90). The topographic height (z) range is as follows, from left to right: Top: 2.3 nm, 2.5 nm, 2.3 nm, 2.4nm; Bottom: 2.8 nm, 3.3 nm, 2.6 nm, 2.5 nm. C) Plots of thickness and width of the sacculi. Mann-Whitney test (****, $P < 0.0001$, $n = 50$) and Mann-Whitney U test (**, $P < 0.0014$) were used for thickness and width data, respectively.Data are representative of three independent AFM images. D) AFM images collected in liquid showing the architecture of *H. pylori* PGN in exponential and late stationary phase. Top: low-resolution images. Bottom AFM high-resolution image of the selected white regions, revealing detailed molecular organization. The two white arrows point toward regions with missing material. The topographic height (z) range is as follows: (top left) 20.4 nm, (bottom left) 11.2 nm, (top right) 26.5 nm, and (bottom right) 19.35. Data are representative of three independent AFM images.

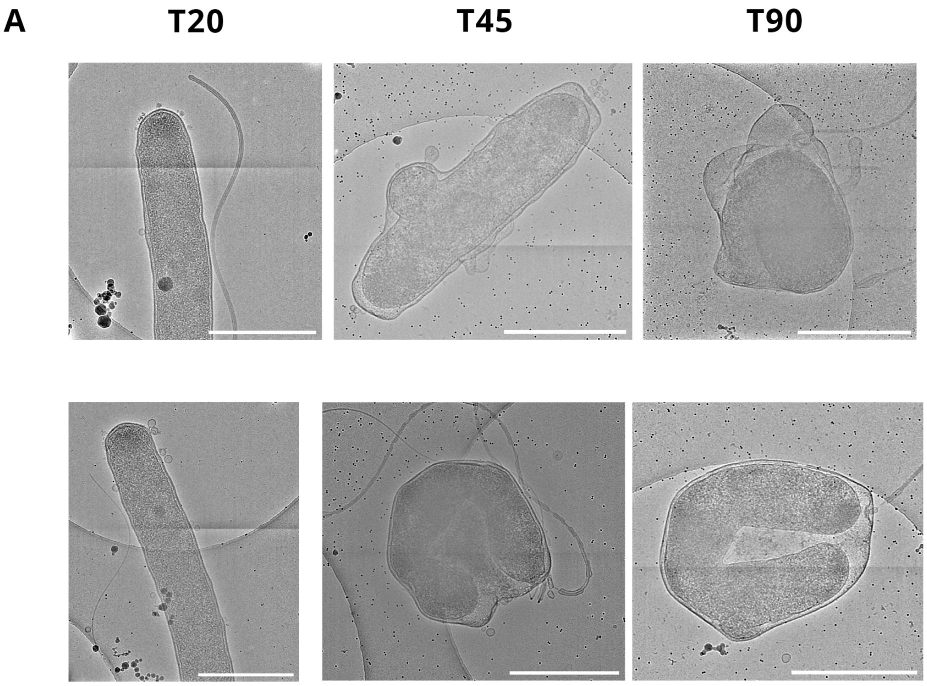


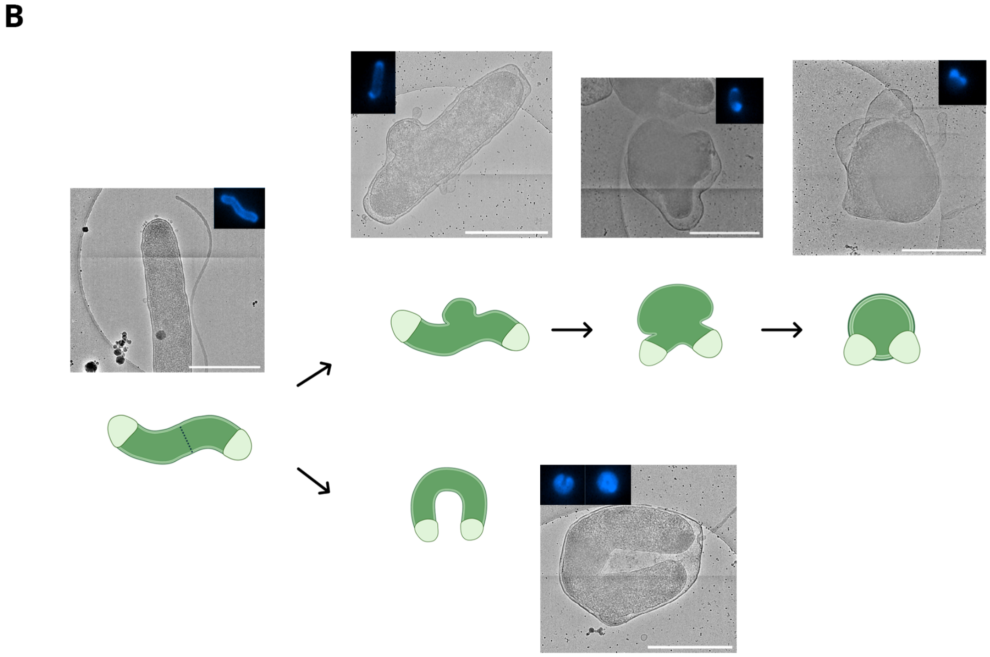


**Fig. S6.** 2D Cryo-EM images of *H. pylori* cells at different stages during the morphological transformation (scale bars: 1μm). A) Bacterial cells were grown in standard conditions and fixed with glutaraldehyde for imaging. Bacteria were captured at the indicated times. B) Cryo-EM images ascribed to stages of the coccoid and U-form transition. HADA fluorescence images typical for each stage are included for comparison.

A

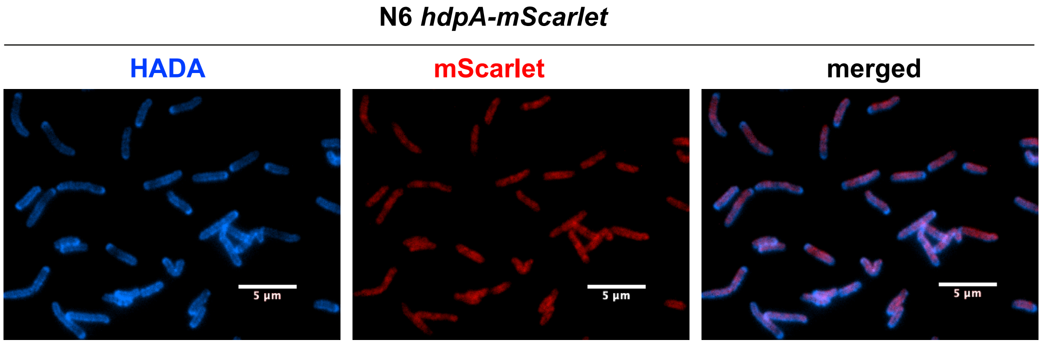


B

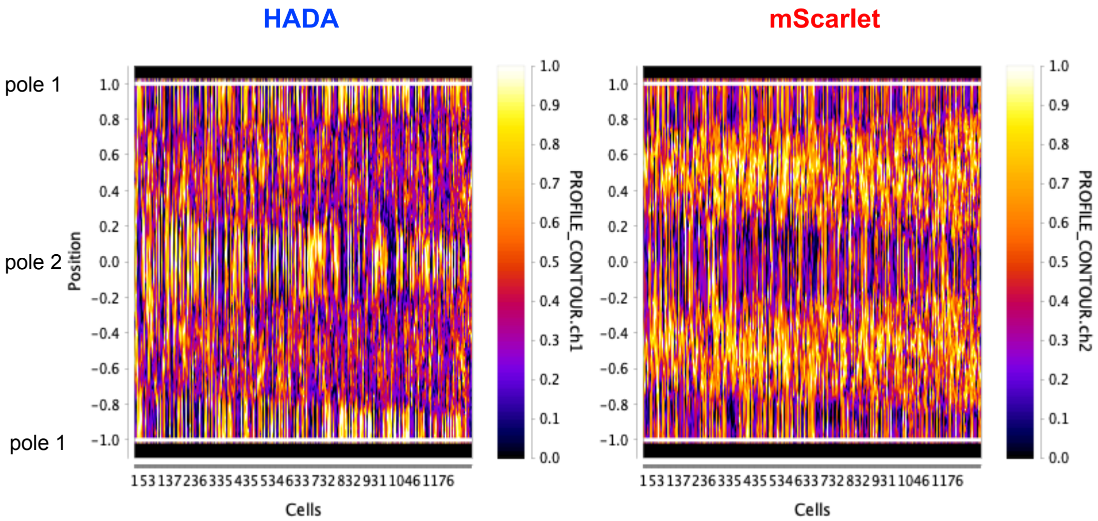


**Fig. S7.** HdpA/Csd3 is involved in the remodeling of the FDAAs in *H. pylori*. A) Fluorescence images of a recombinant strain expressing HdpA-mScarlet (red) imaged at T20 of growth labeled with HADA (blue). (B Demographs of fluorescence intensity of HADA and mScarlet along cell contour sorted by cell length at T20 of growth.

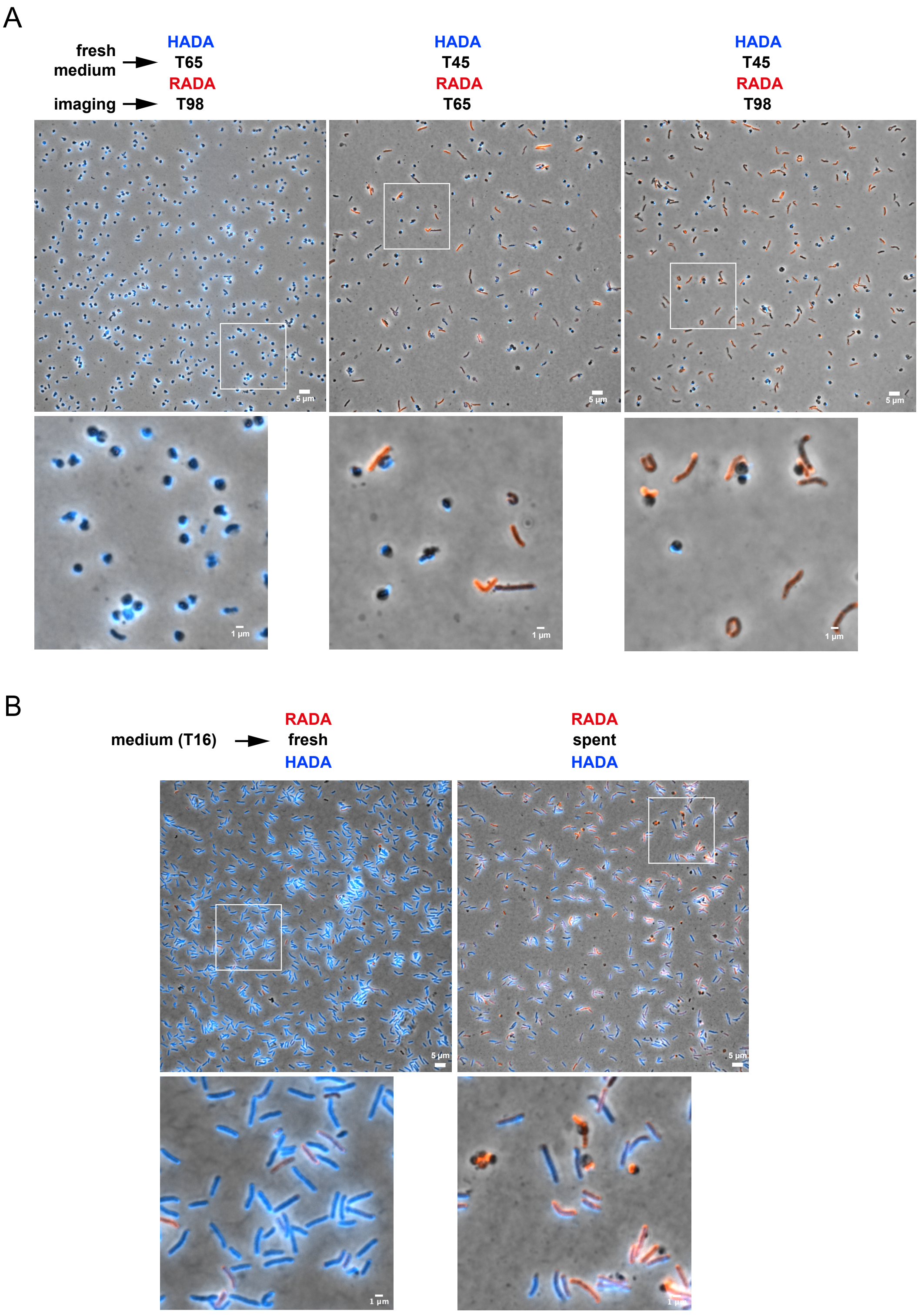


**Fig. S8.** Effects of fresh and spent medium on coccoid formation in *H. pylori*. A) Bacterial cells were cultured in the presence of HADA (H), switched to fresh medium containing RADA (R) and imaged at the indicated times. B) RADA-labeled cells were captured at T16 and transferred into either fresh medium or spent medium from T65 cells, all supplemented with HADA. Cells were imaged at T20.

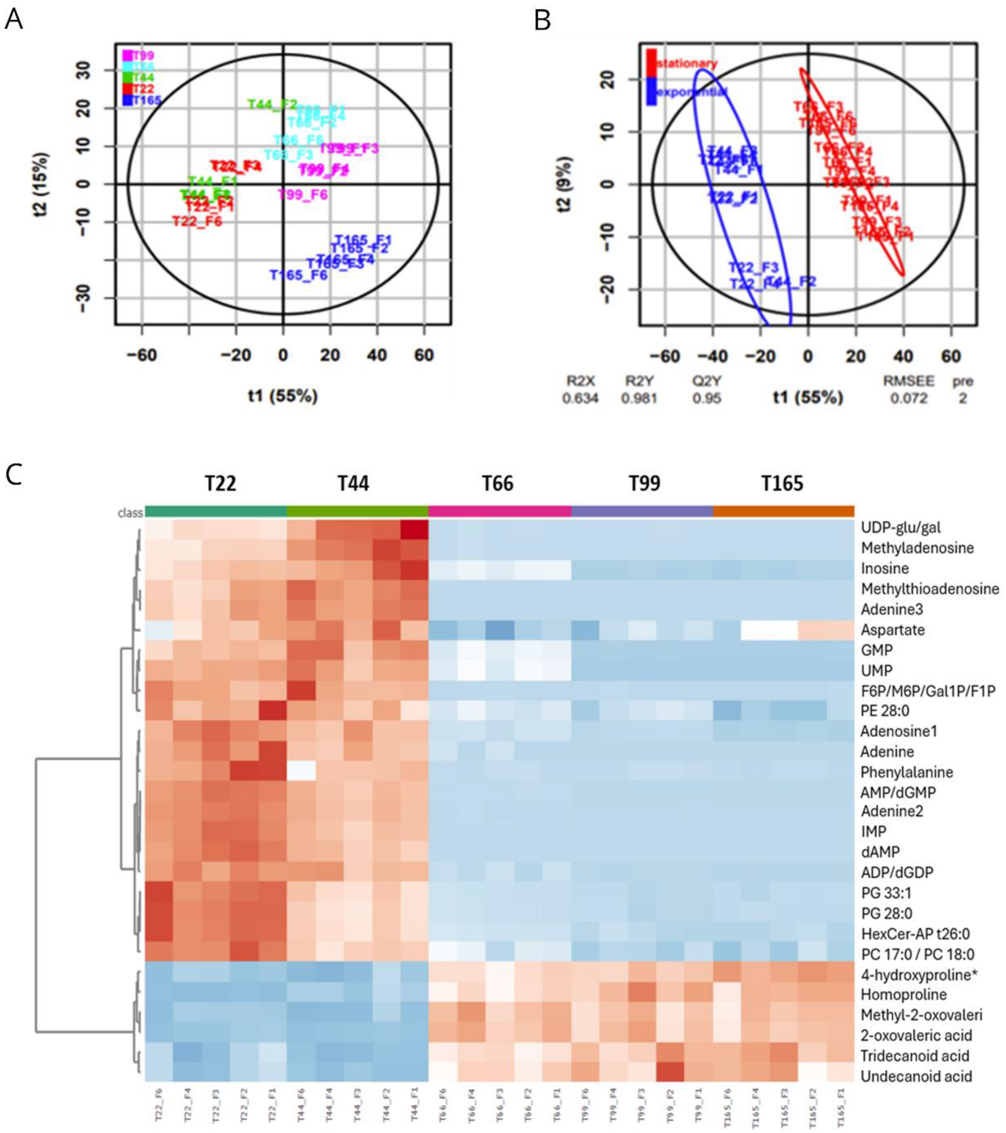


**Fig. S9.** A) PCA score plot of features obtained from LC-MS of *H. pylori* N6 strains at different growth stages. B) PLS-DA score plots show clear separation by PLS component 1 of the exponential phase (blue) and stationary phase (red) of bacteria. C) Heatmap of metabolomic profiles of different growth phases of bacteria based on 22 significantly discriminant metabolites. Data were mean-centered, and unit variance scaled and utilized for heatmap plot. The blue color represents the trend of decrease, red represents a rising trend

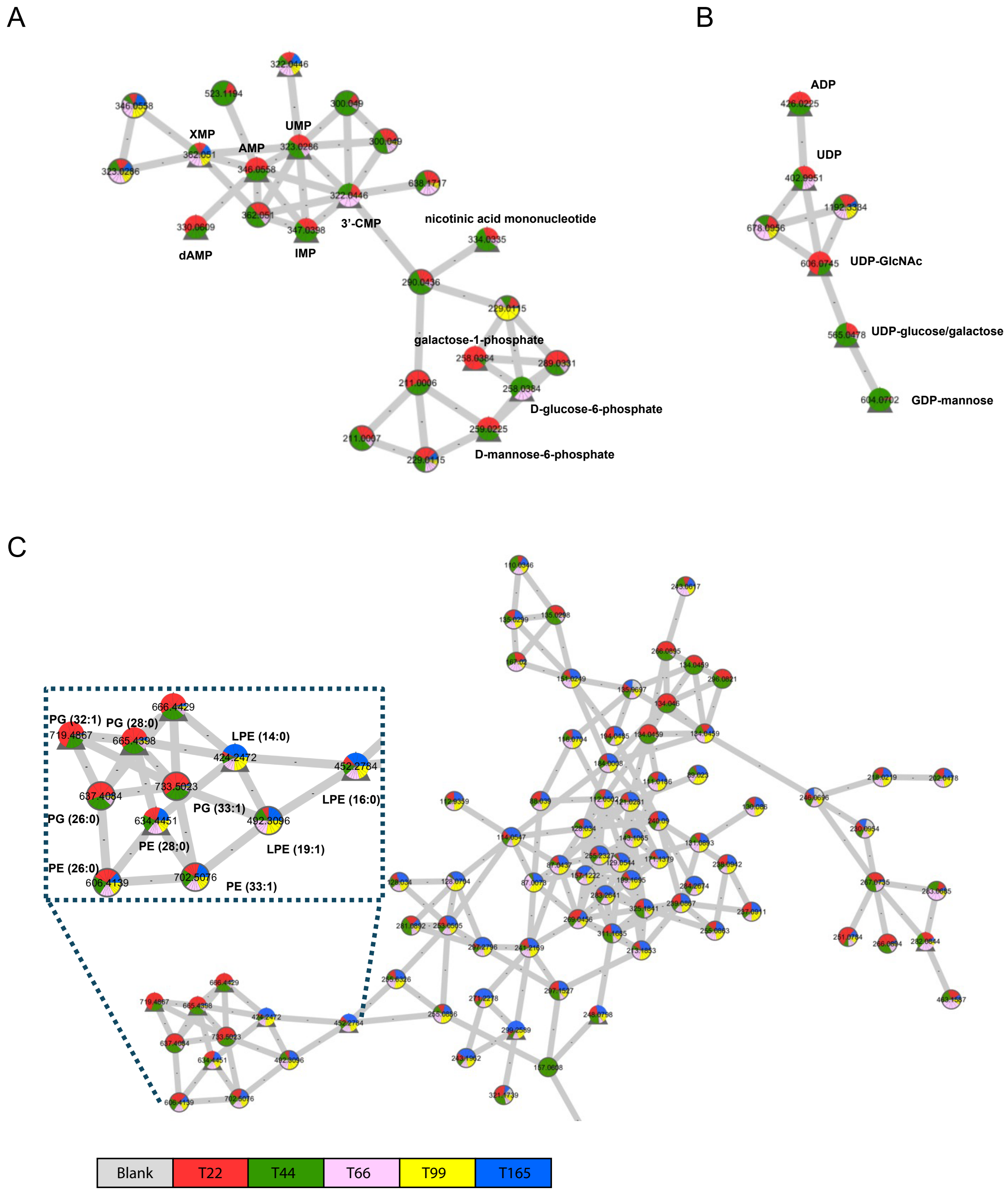


**Fig. S10.** The molecular network was constructed from the crude extracts of *H. pylori* at different growth stages. A) nucleotide cluster  B) hexose-phosphate cluster C) lipid cluster.

**Table S1.** Origin of the 8 strains of *H. pylori* tested in this work

| Strain | Origin | Source | Genome sequence |
|---|---|---|---|
| 26695 | UK | Gastritis patient | available |
| G27 | Italy | Endoscopy biopsy | available |
| B128 | USA | Gastric ulcer patient | available |
| J99 | USA | Duodenal ulcer patient | available |
| N6 | France | Gastritis patient | available |
| P12 | Germany | Duodenal ulcer patient | available |
| SS1 | Australia | Dyspepsia patient | available |
| X47 | USA | Domestic cat with gastritis | available |

**Table S2.** Summary of the four parameters for the 8 strains of *H. pylori*.

| Strain | Rod/Coccoid ratio | Aggregation | Transition period | Plasmid DNA uptake |
|---|---|---|---|---|
| 26695 | 2.9 | Moderate | T40-T135+ | Yes |
| G27 | 2.2 | Mild | T40-T60 | Yes |
| B128 | 2.2 | Severe | T60-T85 | Yes |
| J99 | 1.8 | Mild | T35/45-T85 | Yes |
| N6 | 2.3 | Mild | T40-T85 | Yes |
| P12 | 2.2 | Moderate | T40-T135+ | Yes |
| SS1 | 2.1 | Moderate | T45-T85 | Yes |
| X47 | 2.3 | Moderate (coccoid form) | T65-T110 | No |

**Movie S1-2 (separate files)**
Two examples of time-lapse of the rod to coccoid transition of N6 bacterial cells stained with RADA.
**Movie S3 (separate file).**
Sequential “Z”-slices from a tomogram of a coccoid cell. Because of the large diameter of the N6 coccoid form, the cryo-ET data has significantly reduced signal to noise.
**Movie S4 (separate file).**
Sequential “Z”-slices from a tomogram of a rod-shaped cell.
