## Supplementary figures and images for "Morphological transformation in *Helicobacter pylori* is a dynamic process leading to two types of coccoid"

### Movie S1

#### Slide 1
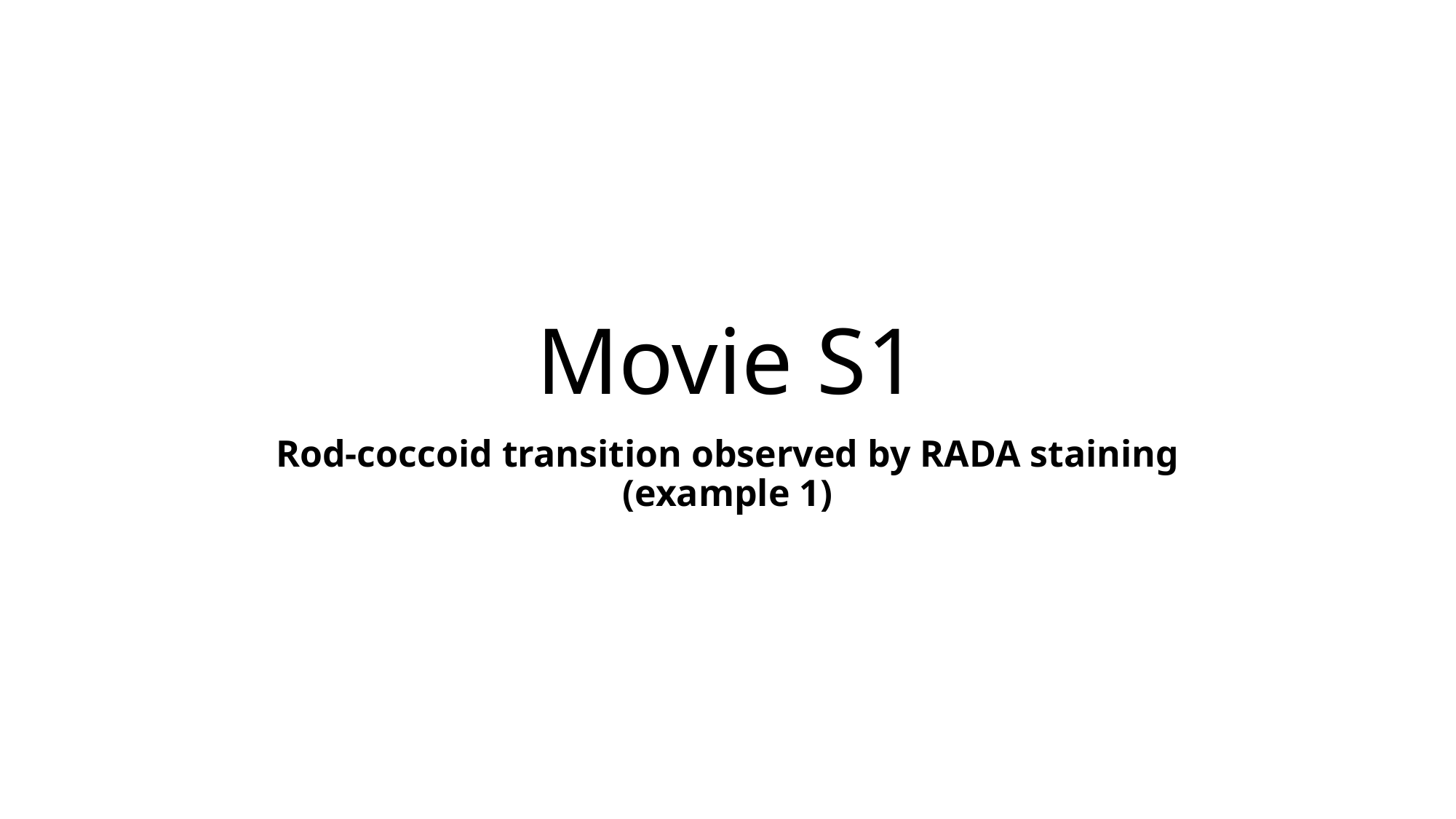

##### Movie S1
Rod-coccoid transition observed by RADA staining (example 1)

#### Slide 2
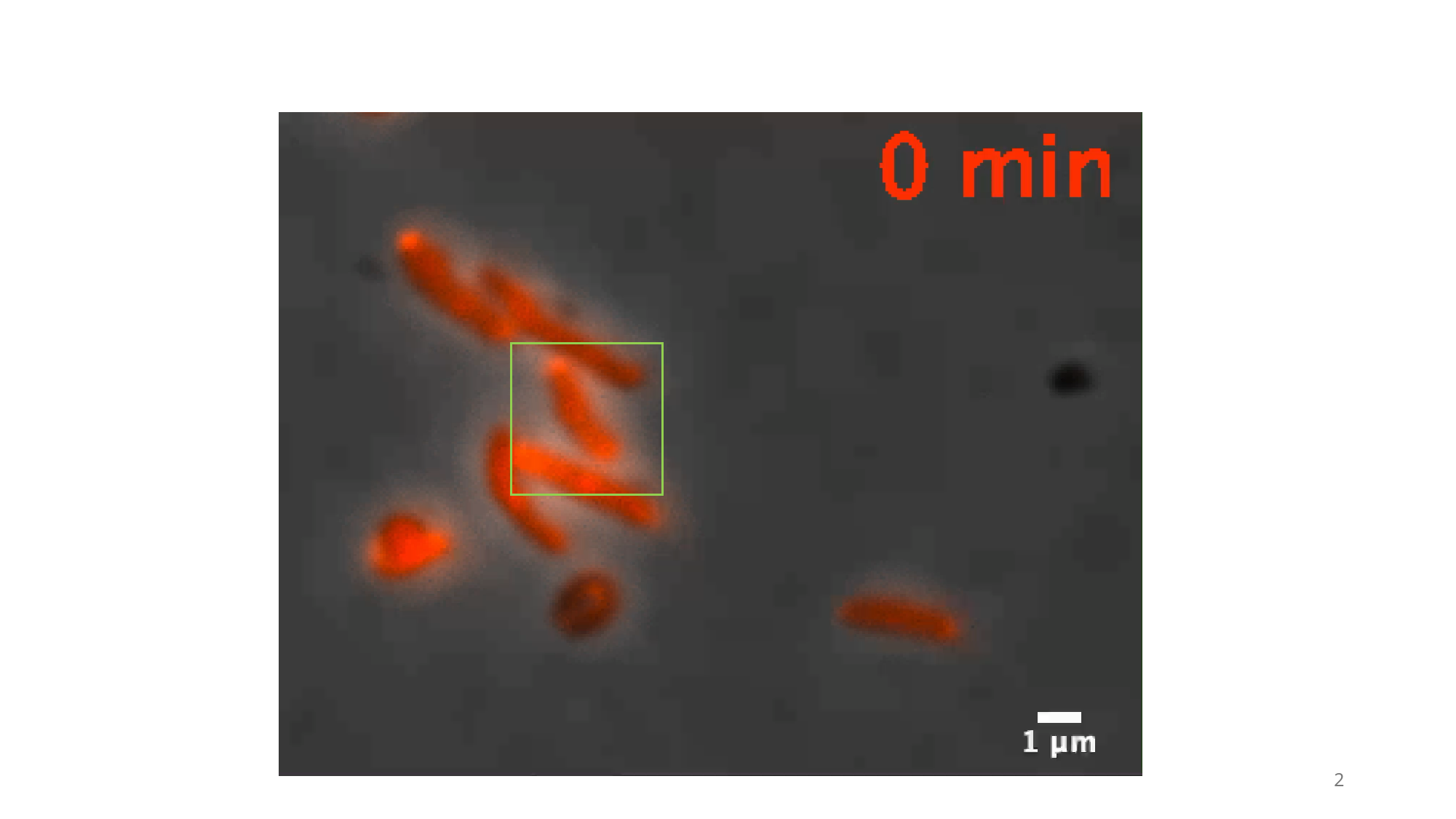

2

### Movie S2

#### Slide 1
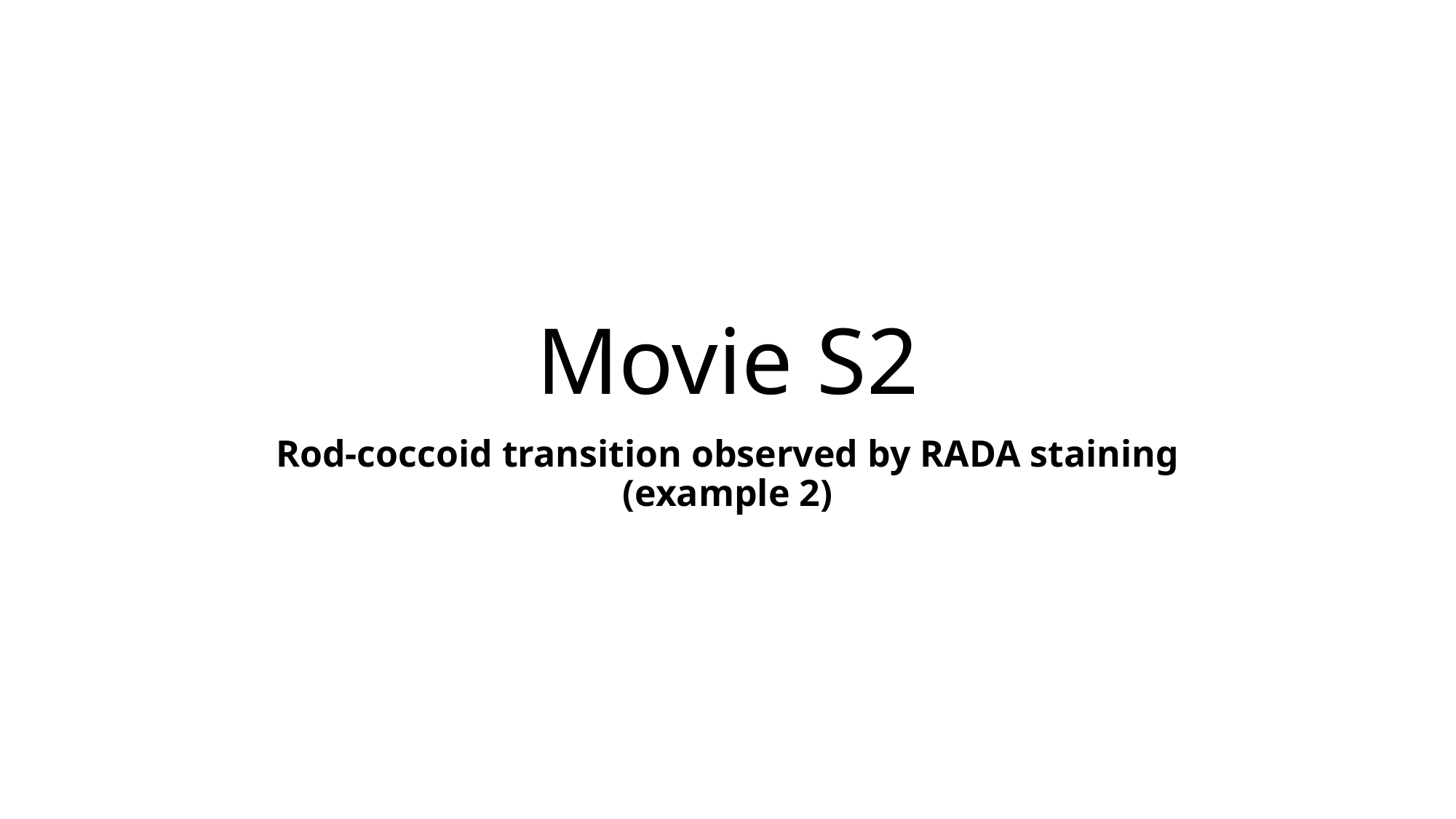

##### Movie S2
Rod-coccoid transition observed by RADA staining (example 2)

#### Slide 2
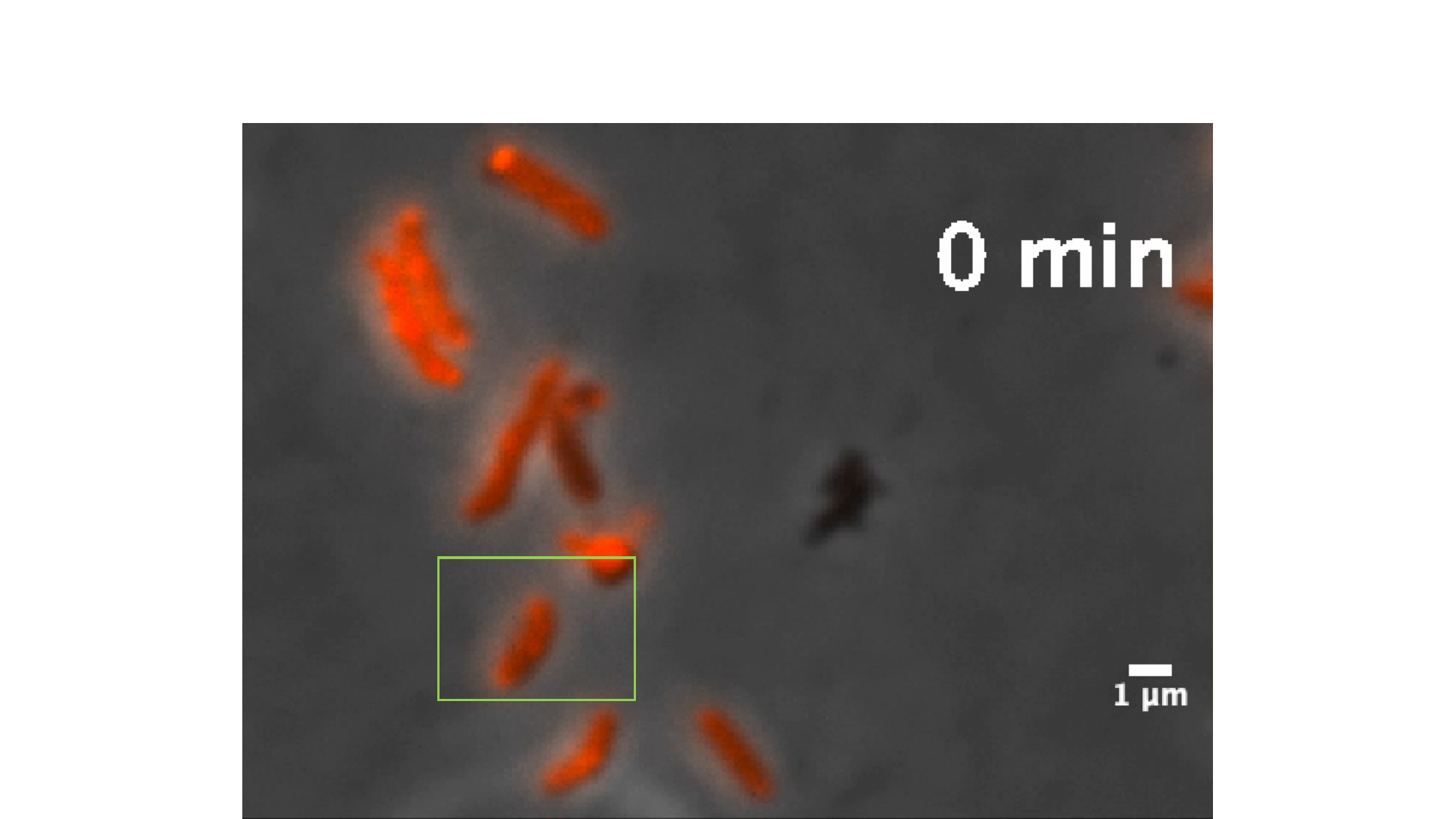

### Movie S3

#### Slide 1
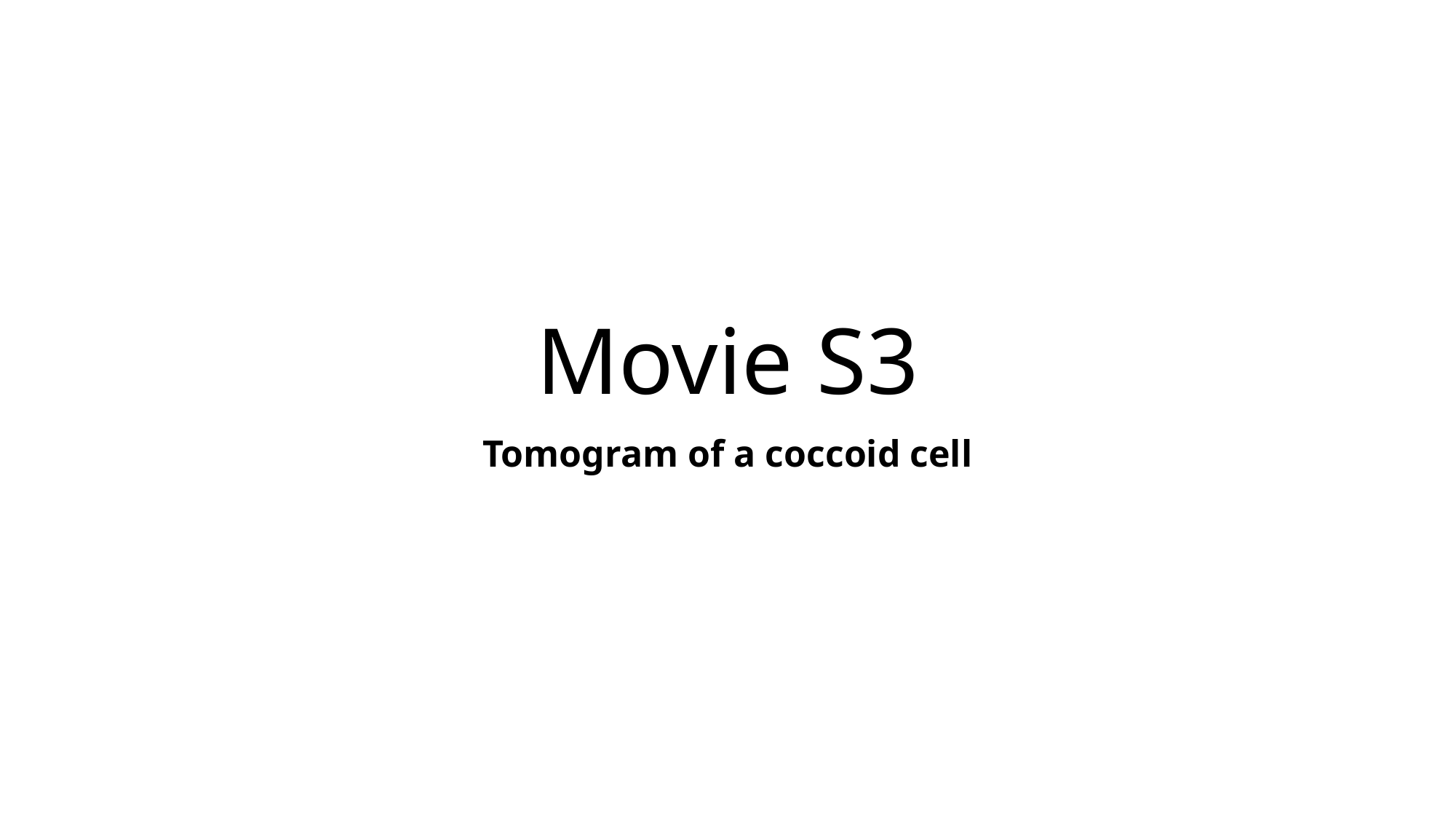

##### Movie S3
Tomogram of a coccoid cell

#### Slide 2
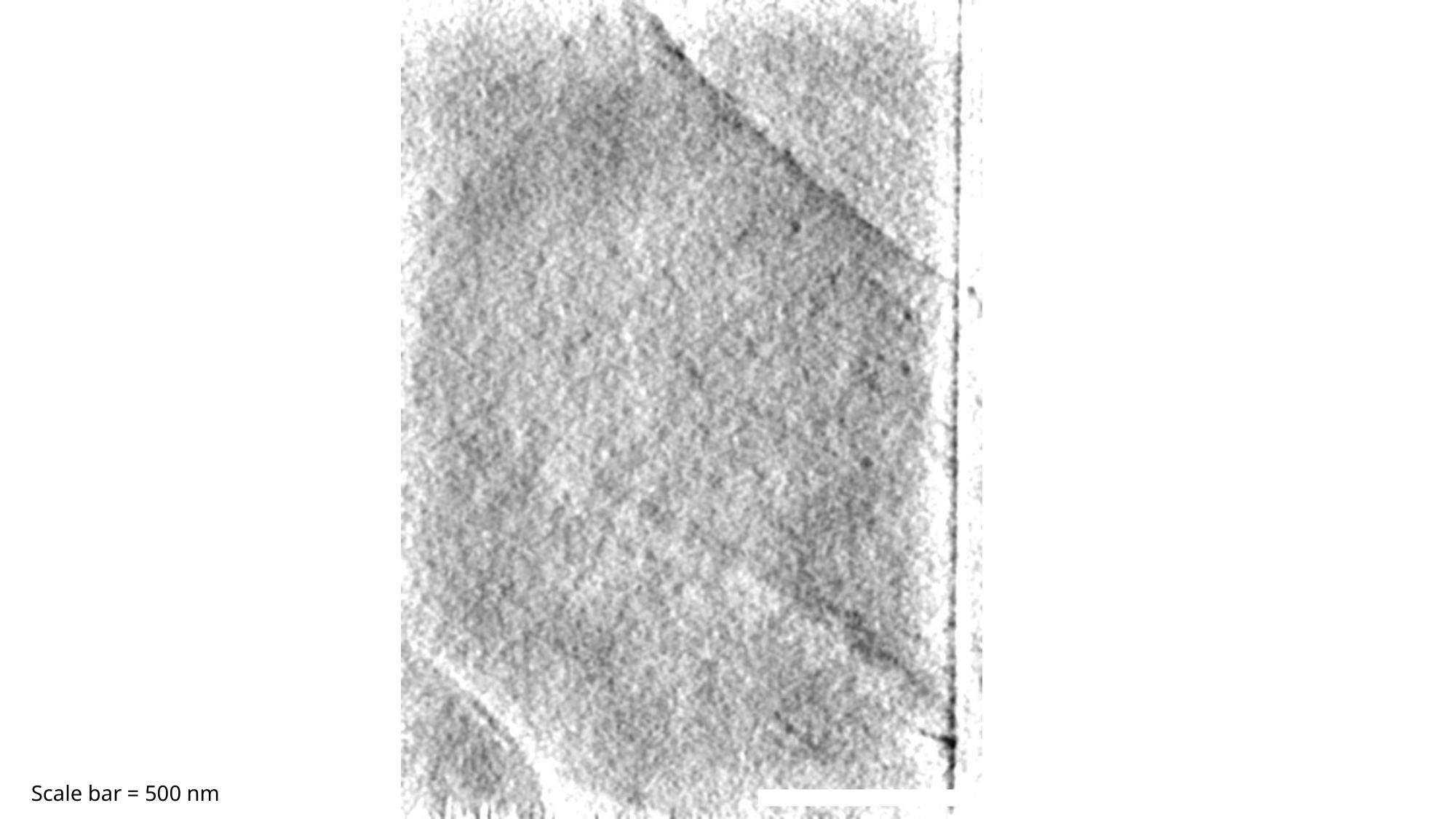

Scale bar = 500 nm

### Movie S4

#### Slide 1
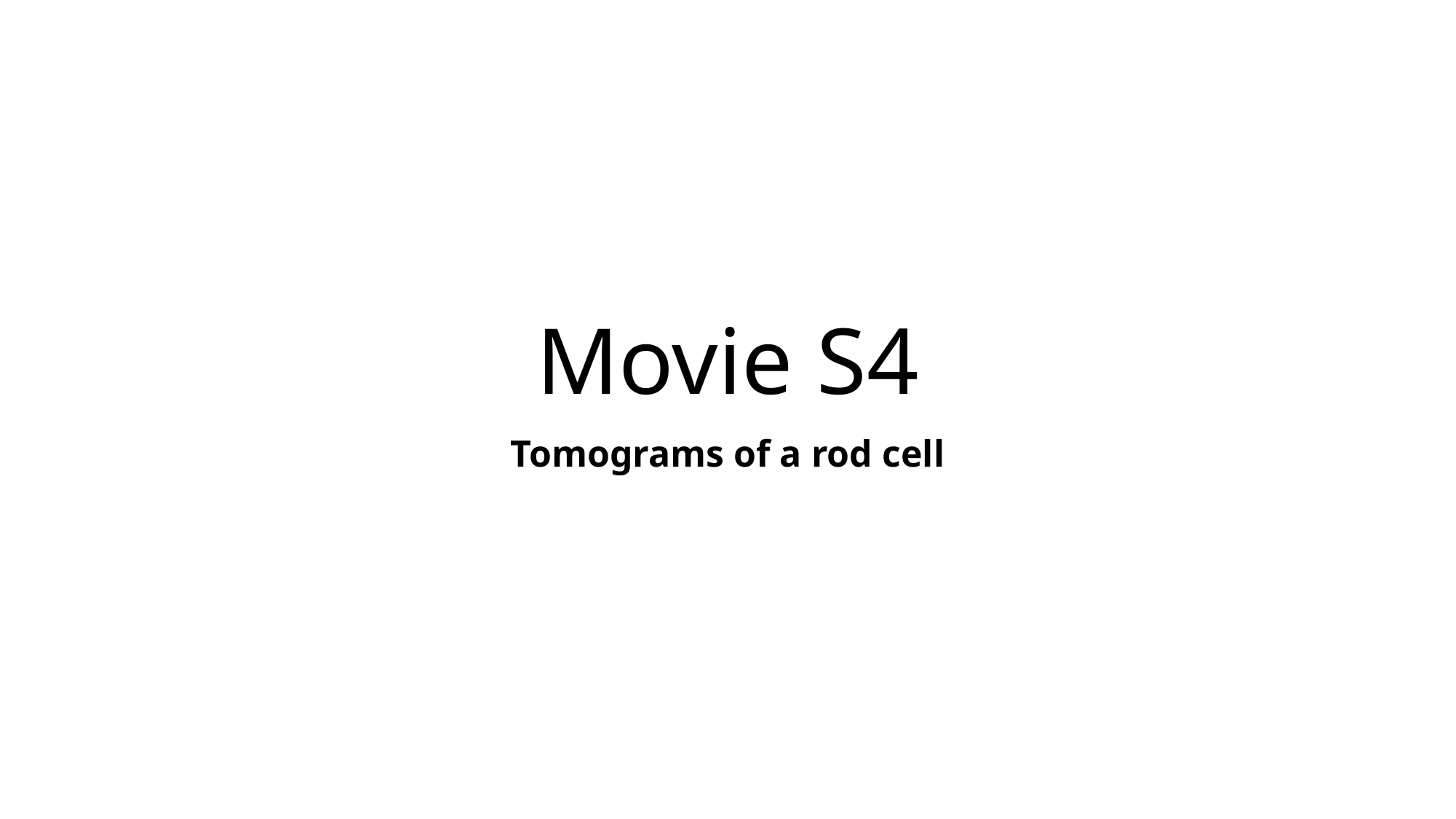

##### Movie S4
Tomograms of a rod cell

#### Slide 2
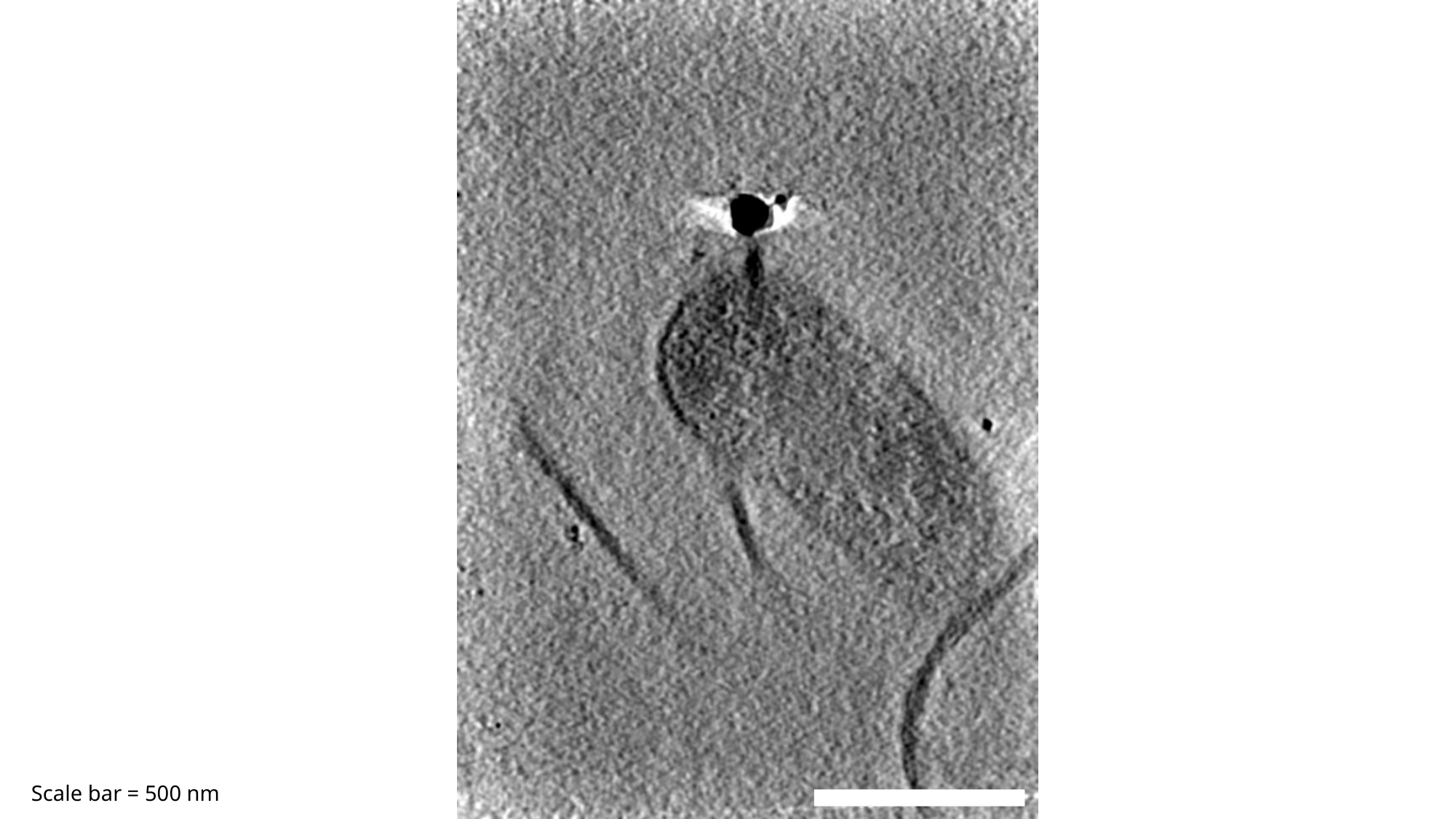

Scale bar = 500 nm
